# Loss of ARID1A accelerates prostate tumourigenesis with a proliferative collagen-poor phenotype through co-operation with AP1 subunit cFos

**DOI:** 10.1101/2024.06.07.597912

**Authors:** Andrew Hartley, Laura C.A. Galbraith, Robin Shaw, Amy Tibbo, Rajan Veeratterapillay, Laura Wilson, Rakesh Heer, Karen Blyth, Hing Leung, Imran Ahmad

## Abstract

Prostate cancer (PC) is the most common male visceral cancer, and second leading cause of cancer mortality in men in the Western world. Using a forward-mutagenesis Sleeping Beauty (SB) transposon-based screen in a Probasin Cre-Recombinase (*Pb-Cre*) *Pten*-deficient mouse model of PC, we identified *Arid1a* loss as a driver in the development of metastatic disease. The insertion of transposon in the *Arid1a* gene resulted in a 60% reduction of *Arid1a* expression, and reduced tumour free survival (*SB:Pten^fl/fl^ Arid1a^INT^* median 226 days vs *SB:Pten^fl/fl^ Arid1a^WT^*293 days, p=0.02),with elevated rates of metastasis (*SB:Pten^fl/fl^ Arid1a^INT^* 75% lung metastasis rate vs 17% *SB:Pten^fl/fl^ Arid1a^WT^,* p<0.001 ). We further generated a *Pb-Cre Pten*- and *Arid1a*-deficient mouse model, in which loss of *Arid1a* demonstrated a profound acceleration in tumorigenesis in *Pten^fl/fl^* mice compared to *Pten* loss alone (*Pb-Cre Pten^fl/fl^Arid1a^+/+^*median survival of 267 days vs Pb-Cre *Pten^fl/fl^ Arid1a^fl/fl^*103 days, p<0.0001). Our data revealed homozygous *Arid1a* loss is required to dramatically accelerate prostate tumourigenesis, resulting in tumours with a less differentiated phenotype and a disorganised stroma. Furthermore, *Arid1a* loss mediated tumour formation in the mouse involved both the anterior and dorsolateral lobes, a unique feature from *Pten*-loss and other reported PC GEMM where tumour formation tends to be limited to the anterior lobes. Analysis of RNA and ChIP -Sequencing data suggests *Arid1a* loss enhanced the function of AP-1 subunit cFos. In clinical PC cohort, ARID1A and cFos levels stratified an aggressive subset of PC with a poor survival outcome with a median of only 30 months.

## Background

Prostate cancer (PC) is the most common cancer in men and the second most common cause of cancer related deaths in the Western world (1). Premature deaths from PC are a result of metastatic and/or recurrent disease for which there are no curative options. Building on our recent research on an *in vivo* Sleeping Beauty (SB) transposon based forward-mutagenesis screen (2, 3), we explore putative role of ARID1A as a mechanism for progressive PC.

Alterations in epigenetic programming are increasingly implicated in the development and progression of cancers. ARID1A is a subunit of the chromatin-remodeling BRM/BRG1-associated factors (BAF) complex, which is a member of the Switch-Induced/Sucrose Non-Fermentable (SWI/SNF) subfamily (4, 5). The BAF complex uses an ATP-dependent chromatin remodeling enzyme, either Brahma (BRM) or Brahma-related gene 1 (BRG1), as the catalytic subunit to remodel chromatin (6). By altering chromatin and nucleosome structures, access to DNA can be altered to epigenetically control gene expression. ARID1A is only found in the BAF complex and is mutually exclusive in the BAF complex with ARID1B, which shares approximately 50% homology (7).

As one of the most mutated epigenetic regulators in cancer (8), ARID1A seems to have a context dependent role in different cancers. ARID1A as part of the BAF complex can mediate chromatin remodeling and gene expression which can be pro- or anti-tumorigenic. Wnt/β-catenin, KRAS, and oestrogen receptor (ER) are all oncogenic pathways which are disrupted when ARID1A is lost (9–11). ARID1A has also been shown to be a tumour suppressor due to its inhibition of cell cycle, mediation of DNA-repair, and high mutation rates in certain cancers such as ovarian cancer (12–15). Similarly, the role of ARID1A in PC remains unclear with both tumour promoting and suppressing effects reported. ARID1A has been shown to regulate oncogenic drivers such as ERG and androgen receptor (16, 17). Our data revealed that homozygous *Arid1a* loss is required to dramatically accelerate prostate tumourigenesis, resulting in tumours with a reduced and disorganised stroma. *Arid1a* loss mediated tumour formation in the mouse involved both the anterior and dorsolateral lobes, a key distinction from *Pten*-loss driven tumours which tend to be limited to the anterior lobes. Finally, the status of PTEN, ARID1A and cFos, as an ARID1A downstream effector, is associated with patient survival outcome.

## Materials and Methods

### Mice

Animal experiments were carried out in line with the Animals (Scientific Procedures) Act 1986 and the EU Directive of 2010 sanctioned by Local Ethical Review Process (University of Glasgow). Mice were maintained on a mixed strain background at the Cancer Research UK Scotland Institute under project licence authority (70/8645 and P5EE22AEE to Professor Hing Leung). Mice were bred and housed in individually ventilated cages under specific pathogen-free conditions on a 12/12-hour light/dark cycle and fed and watered *ad libitum.* Mice were genotyped by Transnetyx using PCR analysis of ear notch tissue.

Alleles used were as follows: *Arr2* Probasin-Cre(18), *Pten*^flox^ (19), T2/Onc3^het-^ (20), *Rosa*^26Lox66SBLox71/+^ (20), and *Arid1a^flox^* (21). Mice were aged until ethically approved clinical endpoints where mice display clinical signs (bladder distension, hunching and/or weight loss), or a palpable prostate tumour >1.2cm. Mice which were culled for reasons other than tumour-associated clinical endpoints were excluded from analysis. Mice were culled when reaching an ageing endpoint of 18 months. All cohort mice were male and were monitored by researchers trained in relevant clinical signs three times per week.

### Cell lines

DU145 (dural metastatic), PC3 (bone metastatic), LNCaP (lymph node metastatic), C4-2 (LNCaP derivative), CWR22 (primary prostate tumour) human prostate cancer cell lines were obtained from ATCC and grown in RPMI-1640 (Sigma Aldrich) and supplemented with 1% L-Glutamine (Gibco) and 10% fetal bovine serum (FBS) (Sigma Aldrich). This medium was used for all cell lines in most instances so will be referred to as standard culture medium (SCM). The 22Rv1 human prostate cancer cell line was grown in RPMI-1640 (Sigma Aldrich) and supplemented with 1% L-Glutamine (Gibco) and with charcoal-stripped FBS (Thermo Fisher) to remove lipophilic materials such as androgen. RWPE human prostate cancer cell line was grown in Keratinocyte serum free medium (Thermo Fisher) supplemented with human recombinant epidermal growth factor (rEGF) and bovine pituitary extract as supplied. Cell cultures were routinely tested for and found to be negative for mycoplasma contamination and were authenticated by the Laboratory of the Government Chemist standards.

Stable *ARID1A* knock out (KO) clones were generated in DU145 cells using a CRISPR/Cas9 plasmid with a specific guide RNA to the *ARID1A* sequence (Santa Cruz, sc-400469) and a homology directed repair plasmid (Santa Cruz, sc-400469-HDR). Amaxa Cell Line Nucleofector Kit (Lonza) was used for electroporating the cells with the plasmids. Setting A023 and nucleofector kit L was used for DU145. Puromycin was used for selection and individual clones were picked following selective pressure. A CRISPR/Cas9 control plasmid with a non-specific guide RNA (Santa Cruz, sc-418922) and an in-house Infra-Red Fluorescent Protein (IRFP) plasmid were used as a control with puromycin as a selectable marker on the IRFP plasmid. As above, puromycin was applied and individual puromycin resistant control clones were selected.

### siRNA treatment

siRNAs were purchased from Dharmacon: ON-TARGETplus Human *ARID1A* siRNA SMARTPool or ON-TARGETplus non-targeting siRNA (Sequences shown in Supplementary Table 1). Cell lines were reverse transfected with siRNAs to a final concentration of 25nM using Lipofectamine RNAiMAX (Invitrogen) following the manufacturer’s protocols with three technical replicates per experiment. To assess siRNA knockdown efficiency, RNA was extracted for quantitative real-time PCR (RT-PCR) analysis.

### Cell growth analysis

Following seeding at 1×10^5^ cells/ml in a 6-well plate, and reverse transfection with siRNAs indicated above, DU145, PC3, and LNCaP cells were counted after 72h. Growth following *ARID1A* knockdown was normalized to the non-targeting control. Each experiment included three technical replicates. For stable DU145 KO clones, cells were seeded and counted after 72h with growth shown relative to empty vector 1. Each experiment included three technical replicates.

### Colony Forming Assay

DU145, PC3, and LNCaP cells were plated at 1×10^5^ cells/ml in a 6-well plate, reverse transfected and incubated with siRNAs for 24h. 200 cells of DU145 and PC3, or 600 cells for LNCaP were then reseeded at a low density in a 10cm dish to allow colonies to form. Cells were then fixed and stained with Crystal Violet (0.5% w/v) and colonies were quantified by fluorescent detection using the Odyssey System (LI-COR).

### Immunoblotting

Immunoblotting performed as previously described(22).Immunoblotting was performed with the following antibodies: ARID1A (Cell Signaling, 12354, 1:1000), ARID1B (Cell Signaling, 92964, 1:1000), AR (Santa Cruz, N-20, 1:1000), PTEN (Cell Signaling, 9559, 1:1000), HSC70 (Santa Cruz Biotechnology, SC-7298), Anti-rabbit IgG, HRP linked antibody (Cell Signaling, 7074, 1:400) and Anti-mouse IgG, HRP linked antibody (Cell Signaling, 7076, 1:400). For all immunoblots images shown are representative of three independent experimental replicates.

### RNA extraction

RNA was extracted from cell lines grown in 6-well plates or from cell pellets using the RNAeasy Mini Kit (Qiagen) as per the manufacturer’s instructions. The optional step to remove genomic DNA using RNase-free DNase (Qiagen) was also included for all samples. RNA was eluted in final step of extraction into RNAse-free molecular grade water and quantified using a Nanodrop (Thermo Fisher). For extraction of RNA from snap frozen tissue, samples were pulverized using a micro-homogenizer, and the resulting powdered tissue was resuspended in RLT-buffer (Qiagen RNeasy Mini Kit) and then further homogenized using Precellys tubes and Precellys Evolution Homogenizer (Bertin Instruments). Once homogenized, RNA was extracted using the RNAeasy Mini Kit (Qiagen) as per the manufacturer’s instructions, including the DNase digestion step.

### Real time – PCR (RT-PCR)

RT-PCR was performed as described previously(3).Briefly, first-strand cDNA was produced by reverse transcription from extracted RNA samples using the High-Capacity cDNA Transcription kit (Applied Biosystems) following the manufacturers protocol. RTPCR was carried out using TaqMan Universal Master Mix (Thermo Fisher Scientific) with primer appropriate Universal ProbeLibrary probes (Roche). Taq-man RTPCR was carried out as previously described(2). The *CASC3* gene was used as the reference to normalise expression levels. Data regarding gene expression is shown relative to levels in control cells. (List of primers and universal probe number are shown in Supplementary Table 1)

### RNA Sequencing

RNA sequencing (RNA-Seq) was carried out as previously described (2). Briefly, the quality of the RNA extracted was tested using an Agilent 220 Tapestation on RNA screentape. Three independent experimental replicates of each sample with three technical replicates were sequenced.

Quality checks and trimming on the raw fastq RNA-Seq data files were performed using FastQC(23), FastP (24) and FastQ Screen(25). RNA-Seq paired-end reads were aligned using HiSat2 version 2.2.1(26) and sorted using Samtools version 1.7(27). Aligned genes were identified using Feature Counts from the SubRead package(28).

Expression levels were determined and statistically analysed using the R environment version (29) and utilizing packages from the Bioconductor data analysis suite(30). Differential gene expression was analysed based on the negative binomial distribution using the DESeq2 package(31) and adaptive shrinkage using Ashr(32).

The reference and annotation genomes Ensembl GRCm 38 (33) was used for the mouse RNA-Seq and ChIP-Seq data and Ensembl GRCh38 (34) was used for the human RNA-Seq data.

Identification of enriched biological functions was achieved using g:Profiler (35), and GSEA version 7.5.1 from the Broad Institute (36).

### Chromatin immunoprecipitation (ChIP) Sequencing

The ChIP assay was performed using the SimpleChIP Enzymatic Chromatin IP Kit with magnetic beads (Cell Signaling Technologies #9003). 25mg of murine prostate tissue from *Pb-Cre Pten^fl/fl^ Arid1a ^+/+^*mice was processed following the manufacturer’s instructions and disaggregated using a Dounce homogeniser. The following antibodies were used: Histone H3 (Cell Signaling, #D2B12, 1:100 dilution), normal rabbit IgG (Cell Signaling, #2727, 1:100 dilution), ARID1A/BAF250A rabbit mAb (Cell Signaling, #12354, 1:100). For immunoprecipitation, samples were incubated with antibodies at 4°C overnight. DNA products were then quantified using Qubit high-sensitivity DNA assay kit (Thermo Fisher, Q32851) and libraries prepared using NEBNext Multiplex Oligos for Illumina Index Primer Set 1 (New England Biolabs, E7335S) and NEBNext Ultra II DNA Library Prep Kit for Illumina (New England Biolabs) library preparation kit. Samples were sequenced on a NextSeq 2000 (Illumina) with 30 million 2×100bp paired end reads by the Glasgow Polyomics next generation sequencing and transcriptomics service.

The consensus peak sets were created using the Nf-Core version 1.2.2(37) of the ChIP-Seq workflow. Transcription Factor binding profiles where obtained from JASPAR 2020(38) and Bedtools (39) was used to identify nearest Transcription Factor to each peak.

Binding Analysis for Regulation of Transcription(40) was used to predict functional transcriptional regulators that bind at cis-regulatory regions to regulate gene expression. Genes with an Irwin Hall p-value below 0.05 were identified by combining the RNA-Seq and ChIP-Seq data.

Further analysis and visualisation was conducted using the R programming language and the Tidyverse(41) set of packages.

Computational analysis was documented at each stage using MultiQC(42), Jupyter Notebooks (43) and R Notebooks(44).

### Human tissue microarray (TMA)

0.5mm^2^ cores of prostate tissue, as identified by pathologists, were removed from a representative area of the formalin-fixed paraffin-embedded (FFPE) block. Tissue was obtained from untreated patients undergoing transurethral prostatectomy (TURP) (repository details from Newcastle REC:2003/11). Patients were diagnosed with PC upon histological examination or by transrectal ultrasound scan (TRUS) between the years of 1988-2005. These samples consisted of clinical T1 (N=33), T2 (N=120), T3 (N=113), and T4 (N=25) stage samples. Only patients who died from PC were included in Kaplan-Meier curves (N=49). Following staining and scoring by the Aperia Imagescope v12.4.6.5003 (Leica Biosystems) of the TMA, scores could be grouped. Scores in the lower or higher interquartile ranges were assigned to ‘low’ or ‘high’ groups respectively Scores that resided in intermediate range were defined as being in the ‘medium’ range.

### Immunohistochemistry (IHC)

IHC staining was performed on 4-μm FFPE sections previously dry heated at 60°C for 2 hours. ARID1A (1:200, 12354, Cell Signaling), Col1a1 (1:200, 93668, Cell Signaling), Ki67 (1:1000, 12202, Cell Signaling), Phospho Serine 473 AKT (1:45, 9271, Cell Signaling) and PTEN (1:70, 9559, Cell Signaling), c-Fos (1:300, ab190289, Abcam), JunD (1:75, sc-271937, Santa Cruz)on the Leica Bond Rx autostainer. Sections treated as previously described(45)

IHC was quantified by using HALO Image Analysis software (Indica Labs). Slides were scanned and analysed using HALO to quantify stain intensity and percentage of cells positive for stain. The Software was trained in each instance to classify and quantify only the stain in the epithelial compartment as only this stain constitutes the tumour. The software then allocated a score to each cell. Histoscore was determined by the following formula: (% cells low intensity) + 2(%cells medium intensity) + 3(%cells high intensity) = Histoscore.

### Statistical analysis

Statistical analyses, except for the RNA-seq and ChIP-Seq datasets, were performed using GraphPad Prism v9.3.1. Testing comprised of unpaired two tailed t-tests, Mann–Whitney, Kaplan–Meier survival analysis and one- and two-way ANOVA with post-tests for multiple comparisons (detailed in figure legends). All experiments were performed in experimental replicates, with technical replicates for each experiment noted. The graphs represent the mean data from the repeated experiment or sample ±SEM.

## Results

### Transposon insertion in the *Arid1a* gene accelerates prostate tumourigenesis and reduces mouse survival

Probasin Cre-recombinase *Pten^flox/flox^* (*Pten^fl/fl^)* mice develop invasive prostate adenocarcinoma and reach clinical endpoint between 9 – 12 months (46). However, these mice rarely develop metastasis and have been aged up to 18 months with the tumours confined to the prostate. We employed forward-mutagenesis Sleeping Beauty transposon based system (2) to generate the *SB:Pten^fl/fl^*(*Pb-Cre4Pten^fl/fl^T2/Onc3^het^Rosa^26Lox66SBLox71/+^*) mouse line, whereby gene expression can be randomly altered to identify novel genetic events that accelerate prostate tumorigenesis. We observed reduced survival among *SB:Pten^fl/fl^* mice when compared to control *Pb-Cre; Pten^fl/fl^* mice (*SB:Pten^fl/fl^*, n=17, median 293 days vs *Pb-Cre;Pten^fl/fll^* mice, n=23, median 469 days (47)) (Figure 1A, left panel). To identify putative driver events, prostate tumours were sequenced, and common transposon insertion sites were identified (2). Four out of twenty-one *SB:Pten^fl/fl^* mice were identified to have insertions in the gene body of *Arid1a (Arid1a^INT^). SB:Pten^fl/fl^ Arid1a^INT^* mice had a significantly reduced survival when compared to *SB:Pten^fl/fl^* mice harbouring insertions affecting other genes (*Arid1a^WT^)* (median 226 days vs 293 days respectively) (Figure 1A, right panel). *Arid1a^INT^* bearing tumours had a 64% reduction in *Arid1a* expression (p<0.0001) (Figure 1B). At clinical endpoint, tumour weights were comparable irrespective of the *Arid1a* status (Figure 1C). Intriguingly, mice habouring tumours with *Arid1a^INT^* were found to have high prevalence of metastatic disease, with all four *Arid1a^INT^*mice developing lymph node metastasis as well as three of the four mice bearing lung metastases (Figure 1D).

**Figure 1:**
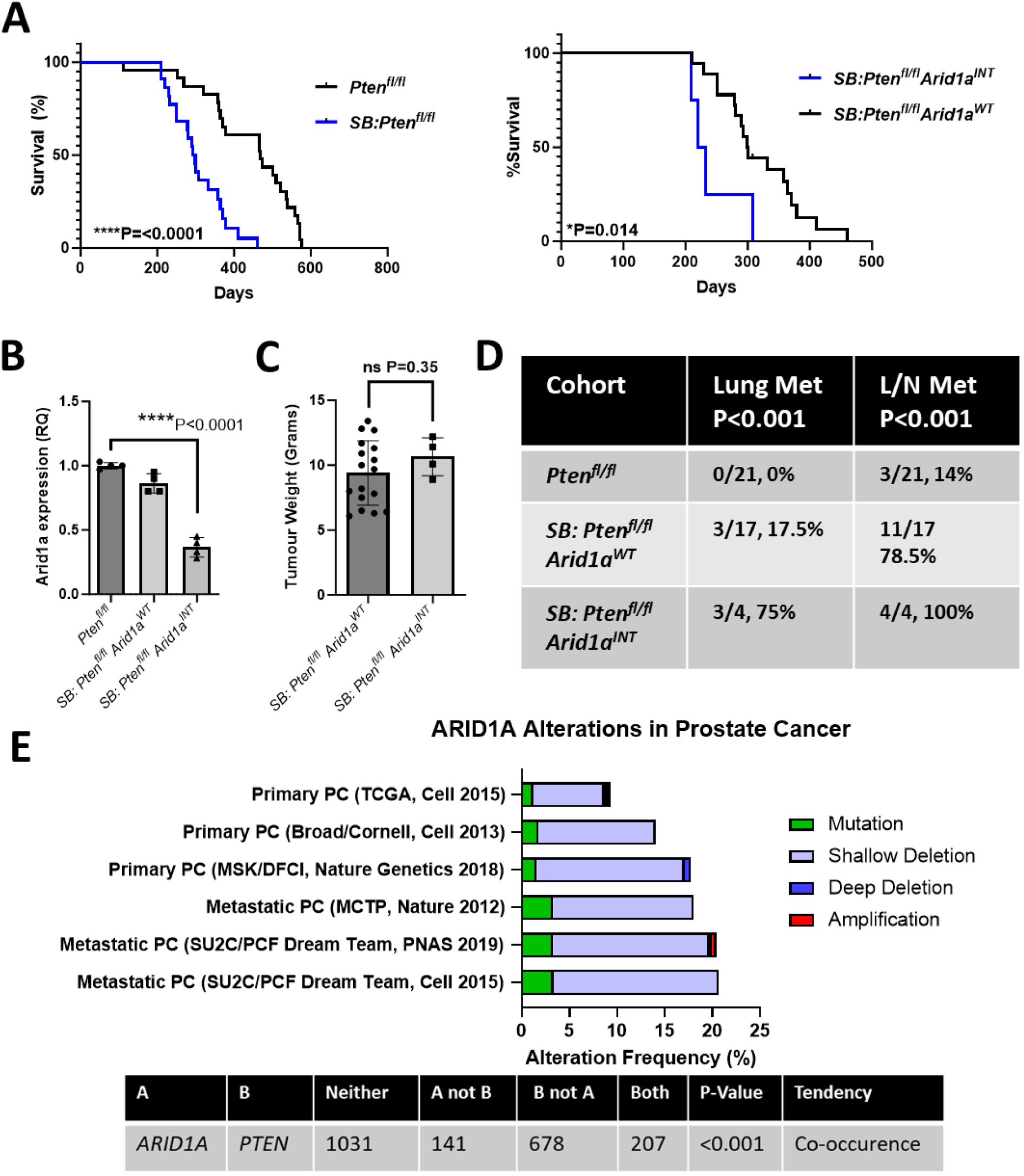
Sleeping Beauty screen identifies *ARID1A* as a candidate driver of advanced prostate cancer. **A.** Kaplan–Meier (log-rank) survival analysis between SB: *Pten^fl/fl^* (n=21) vs *Pten^fl/fl^* (n=23) mice (left panel), and *Pten^fl/fl^Arid1a^WT^* (n=17) vs *Pten^fl/fl^Arid1a^INT^* (n=4) (right panel), ****P=<0.0001; *P=0.014. **B.** RT-PCR for *Arid1a* expression in prostate tumours from *Pten^fl/fl^, SB: Pten^fl/fl^ Arid1a^WT^*, SB: and *SB: Pten^fl/fl^Arid1a^INT^* mice (n=4; each data point represents a different mouse), ****p<0.0001; ANOVA with Tukey’s post-hoc analysis. **C.** Weight in grams of prostate tumours harvested from endpoint tumours of SB: *Pten^fl/fl^ Arid1a^WT^* (n=17) and SB: *Pten^fl/fl^ Arid1a^INT^* (n=4) mice, not significant P=0.35; Mann-Whitney. **D.** Numbers of mice with lung and lymph node (L/N) metastases in *Pten^fl/fl^,* SB: *Pten^fl/fl^ Arid1a^WT^*, and SB: *Pten^fl/fl^ Arid1a^INT^* cohorts with all groups compared by Fisher’s exact test. **E.** *ARID1A* alteration frequency visualised using cBioPortal using the indicated clinical cohorts. Tendency and significance for co-occurrence of *ARID1A* and *PTEN* alterations in these cohorts are also shown.

Using cBioPortal, we visualised *ARID1A* alteration frequencies in multiple (primary and metastatic) PC cohorts (Figure 1E). *ARID1A* was altered between 10-20% of primary PC, and around 20% of metastatic PC (Figure 1E). The common alteration types included shallow deletion and mutation. Importantly, *ARID1A* significantly co-occurred with *PTEN* alterations (Figure 1E), consistent with functional interaction between the two genes in driving prostate tumorigenesis highlighted by our Sleeping Beauty screen.

### Homozygous *Arid1a* deletion drastically accelerates *Pb-Cre;Pten*^fl/fl^ mediatd prostate carcinogenesis

To investigate the functional relevance of *Arid1a* in prostate tumorigenesis *in vivo*, we crossed the *Pb-Cre*;*Pten^fl/fl^*mouse line with *Arid1a^fl/fl^* mice to induce conditional deletion of *Arid1a* and *Pten* in the murine prostate. Homozygous *Pten* loss was functionally confirmed by dramatic upregulated phosphorylation of AKT^Ser473^ (Supplementary Figure 1A), which was maintained regardless of the *Arid1a* status. Similarly, ARID1A loss was confirmed by IHC showing gene copy dependent loss of ARID1A staining in epithelial cells (*Pten^fl/fl^ Arid1a^+/+^*, histoscore of 42; *Pten^fl/fl^ Arid1a^fl/+^,* histoscore of 23.4; *Pten^fl/fl^ Arid1a^fl/fl^*, histoscore of 3) (Supplementary Figure 1B), with ARID1A immunoreactivity also detected in the stroma.

In keeping with our previous findings and the literature, control *Pb-Cre;Pten^fl/fl^ Arid1a^+/+^* mice reached clinical endpoint at a median of 9 months (or 267 days) (46) (Figure 2A). *Pb-Cre;Pten^fl/fl^Arid1a^fl/+^*mice had similar survival outcomes to the control *Pb-Cre*;*Pten^fl/fl^ Arid1a^+/+^* mice: *Pten^fl/fl^Arid1a^fl/+^* (n=19) median 236 days vs. *Pten^fl/fl^ Arid1a^+/+^* (n=10) median 267 days, p=0.83 (Figure 2A). *Pb-Cre;Pten^fl/fl^Arid1a^fl/fl^*mice however developed prostate tumours rapidly, leading to a significant reduction in their survival compared to controls: *Pten^fl/fl^Arid1a^fl/fl^* (n=8) median 103 days vs. *Pten^fl/fl^ Arid1a^+/+^* (n=10) median 267 days, p<0.0001 (Figure 2A). Tumour weights at endpoint were comparable among all three genotypes (*Pten^fl/fl^ Arid1a^+/+^* mean 0.72g, *Pten^fl/fl^Arid1a^fl/+^* mean 0.63g, *Pten^fl/fl^Arid1a^fl/fl^*mean 0.71g), signifying the rapid nature of prostate tumorigenesis in *Pb-Cre;Pten^fl/fl^Arid1a^fl/fl^* mice (Figure 2A, 2B).

**Figure 2:**
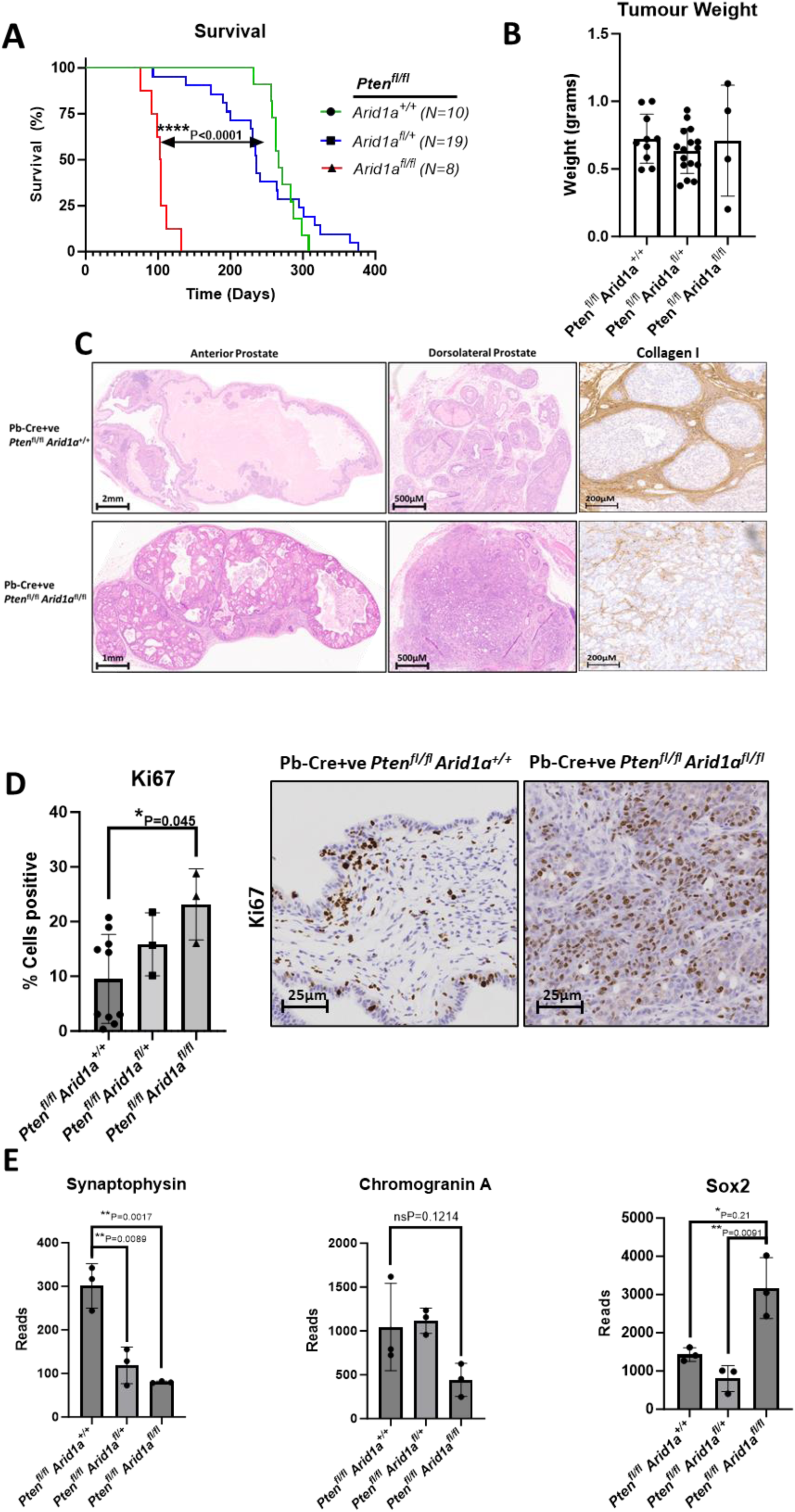
Loss of *Arid1a* in a *Pb-Cre;Pten^fl/fl^* mouse drives aggressive and invasive carcinoma of dorsolateral prostate. **A.** Kaplan–Meier (log-rank) curve demonstrating survival of *Pb-Cre;Pten^fl/fl^Arid1a^+/+^* controls *(n=10), Pb-Cre;Pten^fl/fl^Arid1a^fl/+^ (n=19) and, Pb-Cre;Pten^fl/fl^Arid1a^fl/fl^* (n=8) mice, ****P<0.0001; log-rank (Mantel-Cox) test. **B.** Weight in grams of prostate tumours harvested from endpoint tumours of *Pb-Cre;Pten^fl/fl^Arid1a^+/+^ (n=10), Pb-Cre;Pten^fl/fl^Arid1a^fl/+^ (n=16), Pb-Cre;Pten^fl/fl^Arid1a^fl/fl^* (n=4, a subset of those shown in A where tissue was collected and weight recorded) genotypes. Each data point represents and individual mouse. **C.** Representative endpoint dorsolateral prostate lobes of *Pb-Cre;Pten^fl/fl^Arid1a^+/+^* and *Pb-Cre;Pten^fl/fl^Arid1a^fl/fl^* tumours stained by IHC for Collagen I. **D.** Representative staining for Ki67, and quantification of Ki67-positive total epithelial cells of *Pb-Cre;Pten^fl/fl^Arid1a^+/+^ (n=10), Pb-Cre;Pten^fl/fl^Arid1a^fl/+^ (n=3), Pb-Cre;Pten^fl/fl^Arid1a^fl/fl^* (n=3), *P=0.045, ANOVA with Tukey’s analysis. Each data point represents and individual mouse and a subset of those in A. **E.** RT-PCR for expression of genes of interest in prostate tumours from respectively genotypes (n=3; each data point represents a different mouse).

Homozygous loss of *Arid1a* resulted in epithelial-dense tumours involving both anterior and dorsolateral lobes (Figure 2C), contrasting to tumour formation in the control *Pb-Cre;Pten^fl/fl^ Arid1a^+/+^* mice being limited to the anterior lobes. Furthermore, tumours from *Pb-Cre;Pten^fl/fl^Arid1a^fl/fl^* mice exhibited a distinct morphology with marked loss of a structured stromal compartment as illustrated by collagen I staining (Figure 2C). In keeping with a role for *Arid1a* in tumour morphology, loss of *Arid1a* also led to a reduction in luminal marker keratin 8 (*Pb-Cre;Pten^fl/fl^ Arid1a^+/+^*, histoscore of 77.3; *Pb-Cre;Pten^fl/fl^ Arid1a^fl/+^,* histoscore of 56.8; *Pb-Cre;Pten^fl/fl^ Arid1a^fl/fl^*, histoscore of 43.4) and an elevation in basal marker keratin 5 (*Pb-Cre*;*Pten^fl/fl^ Arid1a^+/+^*, histoscore of 18.8; *Pb-Cre;Pten^fl/fl^ Arid1a^fl/+^,* histoscore of 21.7; *Pb-Cre*;*Pten^fl/fl^ Arid1a^fl/fl^*, histoscore of 62.7) (Supplementary Figure 2). Progressive deletion of *Arid1a* loss was also associated with a more proliferative phenotype with gene copy dependent elevation in Ki67-positive cell staining: *Pb-Cre;Pten^fl/fl^ Arid1a^+/+^*, 9.5% cells positive; *Pb-Cre;Pten^fl/fl^ Arid1a^fl/+^,* 15.8% cells positive; *Pten^fl/fl^ Arid1a^fl/fl^*, 23.2% cells positive (Figure 2D). Loss of *Arid1a* promoted prostate tumorigenesis, with rapid tumour formation and altered tumour morphology (less differentiated epithelial compartment and reduced/disorganised stroma). Furthermore, upregulated *Sox2* mRNA expression in *Pten^fl/fl^ Arid1a^fl/fl^*tumours is consistent with a less differentiated phenotype, while we found no evidence of neuro-endocrine differentiation with reduced *SYP* (Synaptophysin) and equivocal *CHGA* (Chromogranin A) mRNA expression (Figure 2E). Collectively, our findings support our hypothesis that loss of *Arid1a* cooperates with *Pten* loss in *in vivo* prostate tumorigenesis as suggested by the Sleeping Beauty screen (Figure 1).

To characterise the relationship of *Arid1a* loss with wildtype or heterozygous loss of *Pten* in prostate tumorigenesis, we generated the *Pb-Cre; Arid1a^fl/fl^* (*Pten^+/+^*or *Pten^fl/+^*) mouse cohorts. Previous work has already demonstrated that *Pb-Cre;Pten^fl/+^* mice do not develop adenocarcinoma of the prostate (46). *Pb-Cre;Pten^+/+^ Arid1a^fl/fl^* mice did not develop any prostate tumours (Supplementary Figure 3A), although there was evidence of prostate intraepithelial neoplasm (PIN), predominantly in the dorsolateral lobe (Supplementary Table 2A). In contrast, the *Pb-Cre;Pten^fl/+^ Arid1a^fl/fl^* mouse cohort developed tumours in 5 of 12 mice, with three reaching clinical endpoint, along with PIN formation (Supplementary Figure 3A, Supplementary Table 2A). The clinical endpoint tumours of the *Pb-Cre;Pten^fl/+^ Arid1a^fl/fl^* mice were morphologically similar to *Pb-Cre;Pten^fl/fl^Arid1a^fl/fl^* tumours, with a dense tumour texture (Supplementary Figure 3C showing bladder distension due to tumour growth). Given these striking similarities, we hypothesised that inactivation of the remaining *Pten* allele contributes to tumour formation in *Pb-Cre;Pten^fl/+^ Arid1a^fl/fl^* mice, and studied PTEN immunoreactivity in clinical endpoint *Pb-Cre;Pten^fl/+^ Arid1a^fl/fl^* tumour (Supplementary Figure 3D). Indeed, we observed reduced PTEN protein levels in the endpoint *Pb-Cre;Pten^fl/+^ Arid1a^fl/fl^* tumours, while PTEN protein remained intact in benign glands without tumour formation. Hence, *de novo* inactivation of the remaining *Pten* allele may functionally replicate a *Pb-Cre;Pten^fl/fl^Arid1a^fl/fl^*genotype in driving tumorigenesis. Besides differences in tumour morphology, *Arid1a* mediated tumour formation was noted to affect both the anterior and dorsolateral lobes, a key distinction from *Pten*-lost driven tumours which tend to be limited to the anterior lobes (while PIN formation was evident even in the dorsolateral lobes following homozygous *Pten* deletion alone, Supplementary Table 2B).

### Transcriptomic analysis of combined *Pten*- and *Arid1a-* deficient tumours

With a background of homozygous *Pten* deletion, homozygous *Arid1a* loss resulted in substantially more altered gene expression when compared to heterozygous *Arid1a* loss, namely 1540 and 183 genes respectively, with only 132 shared genes (Figure 3A). Principal component analysis (PCA) showed the largest variance compared to control is only achieved following homozygous loss of *Arid1a* while heterozygous loss closely clusters with controls (Figure 3B). Of note, the three tumours from *Pb-Cre;Pten^fl/fl^Arid1a^fl/+^*mice exhibited substantial heterogeneity at the transcriptome level. This is reminiscent of the heterogenous endpoints observed in the *Pb-Cre;Pten^fl/fl^Arid1a^fl/+^* mice, with some mice reaching clinical endpoint as early as 76 days while others as late as 130 days (Figure 2A). *Pb-Cre;Pten^fl/fl^Arid1a^fl/+^*tumours only had 183 significantly dysregulated genes compared to *Pb-Cre;Pten^fl/fl^*, indicating a single copy loss of *Arid1a* does not cause large transcriptional changes (Figure 3C, top panel). In contrast, of the 1540 significantly dysregulated genes following homozygous *Arid1a* loss, with 1143 genes downregulated and only 397 genes upregulated (Figure 3C, bottom panel). The observation that nearly 3 fold more genes were downregulated than upregulated following homozygous *Arid1a* loss is consistent with the notion that *Arid1a* more frequently opens chromatin than closes it (48). Five of ten upregulated cell signalling networks in the *Pb-Cre;Pten^fl/fl^ Arid1a^fl/fl^* tumours were related to cell cycle control (Figure 3D). Geneset enrichment analysis (GSEA) identified the enriched phase of cell cycle signalling to be around the G2/M phase transition, with key regulators of this checkpoint enriched including AURKA, PLK1, NEK2, CCNA2 (Figure 3E).

**Figure 3:**
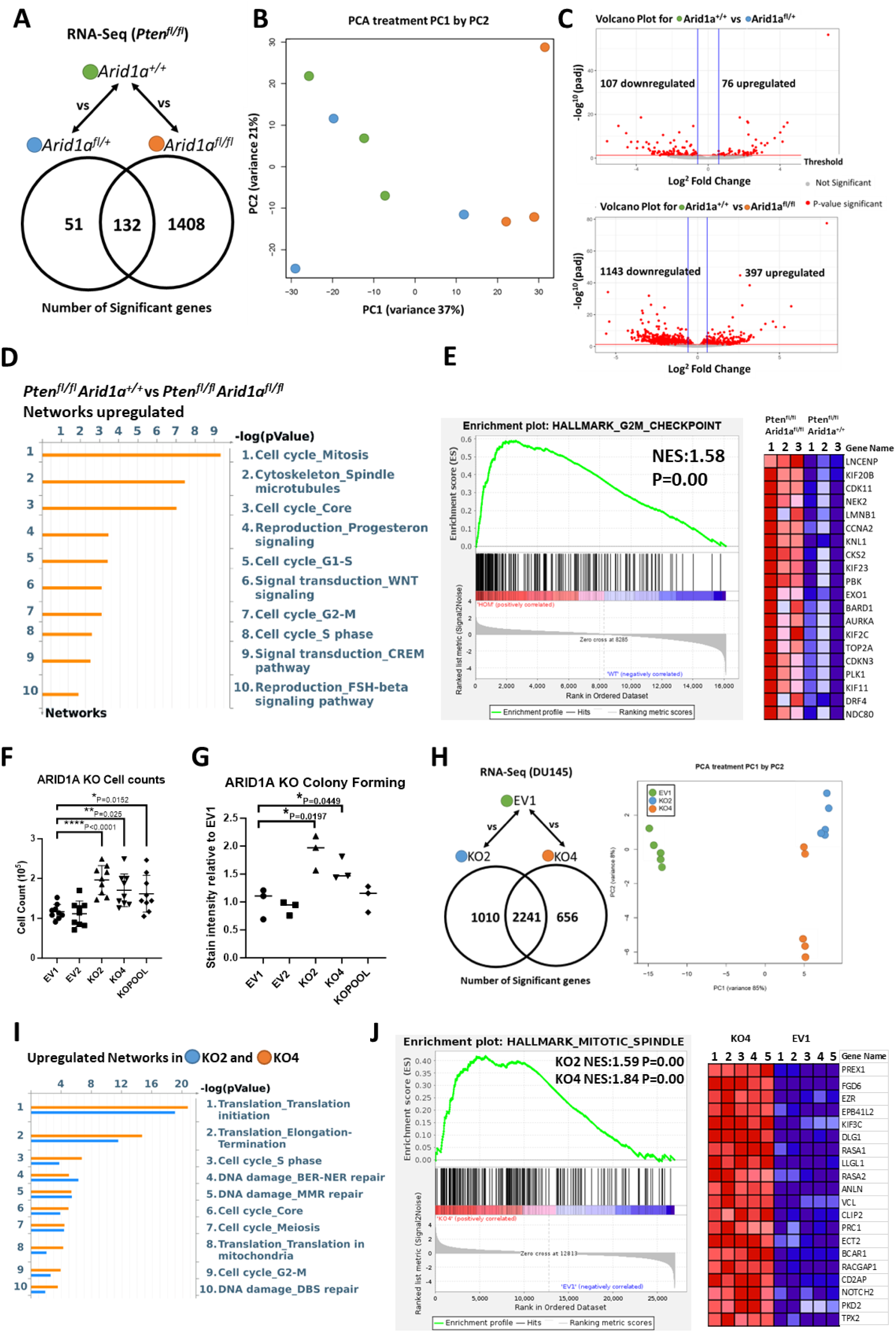
Loss of ARID1A elevates cell cycle signalling in Pten-deficient Tumours. **A.** Number of significant genes (P<0.05, Fold Change >1.5) from RNA-Seq analysis of endpoint prostate tumours comparing *Pb-Cre;Pten^fl/fl^ Arid1a^+/+^* (n=3) compared to *Pb-Cre;Pten^fl/fl^ Arid1a^fl/+^* (n=3) or *Pb-cre;Pten^fl/fl^ Arid1a^fl/fl^* (n=3) mice. **B.** Principal component analysis (PCA) showing comparison and variance of individual mouse samples of indicated cohorts. **C.** Volcano plot showing up and downregulated genes (P<0.05, Fold Change >1.5) in *Pten^fl/fl^ Arid1a^fl/+^* (n=3) and *Pten^fl/fl^ Arid1a^fl/fl^* (n=3) cohorts. **D.** Significantly upregulated cell signalling networks visualised using Metacore in *Pb-Cre;Pten^fl/fl^ Arid1a^fl/fl^ compared* to *Pb-Cre;Pten^fl/fl^ Arid1a^+/+^* tumours. **E.** Gene set enrichment analysis showing 1.58 normalised enrichment score (NES) in Hallmark G2M checkpoint from *Pb-Cre*;*Pten^fl/fl^ Arid1a^fl/fl^* mice. Most significant genes of enrichment shown in heatmap with z-score indicated between +2 to −2 with colour gradient of red to blue. **F.** Fold change in cell count for DU145 EV clones compared to *ARID1A* KO clones after 72 h of growth, *P=0.015, **P=0.0025, ****P<0.0001; ANOVA with Tukey’s analysis. Each data point represents a single technical replicate, three of which made up each experimental replicate, error bars showing SE. **G.** Stain intensity of colony growth from colony forming assay. Relative growth relative to EV1. *P=0.02 EV1 vs KO2, *P=0.045 EV1 vs KO4, ANOVA with Tukey’s analysis. Each point represents an experimental replicate each made up of three technical replicates, error bars showing SEM. **H.** Number of significant genes (P<0.05, Fold Change >1.5) from RNA-Seq analysis comparing EV1 (n=5) compared to KO2 (n=5) or KO4 (n=5). Principle component analysis (PCA) showing comparison and variance of individual samples of indicated cell clones. **I.** Significantly upregulated cell signalling networks visualised using Metacore in KO2 and KO4 compared to EV1. **J.** Gene set enrichment analysis showing 1.59 (KO2) and 1.84 (KO4) normalised enrichment score (NES) in Hallmark Mitotic Spindle compared to EV1. Genes from leading edge of enrichment shown in heatmap with z-score indicated between +2 to −2 with colour gradient of red to blue.

Consistent with data from analysis of *in vivo* tumours, knockout of ARID1A in the human prostate cancer DU145 cells significantly promoted growth, increasing cell counts in DU145 ARID1A knockout KO2, KO4 clones and KO pool cells by 68%, 45% and 38% respectively (Supplementary Figure 4, Figure 3F). Likewise, colony forming capabilities were elevated by 90% in KO2 and 57% in KO4 cells, though not in the KO Pool cells (Figure 3G). We further carried out transcriptomic analysis on KO2 and KO4 cells, comparing to DU145 empty vector (EV1) control cells. PCA confirmed close similarities among the ARID1A KO cell clones when compared to empty vector controls (Figure 3H). Network analysis identified that knockout of ARID1A upregulates cell cycle pathways, as well as increasing translation, suggesting global changes in growth (Figure 3I). GSEA also validated this effect on cell cycle, identifying an elevation in mitotic spindle formation in DU145 KO clones (Figure 3J).

### Loss of ARID1A correlates with upregulation of AP-1 subunit cFos and identifies patients with reduced survival

To gain molecular insight into *Arid1a*-mediated epigenetic changes in prostate tumorigenesis, we performed Chromatin Immunoprecipitation Sequencing (ChIP-Seq) on *Pb-Cre;Pten^fl/fl^* tumours to interpret data from transcriptomic analysis. Binding Analysis for Regulation of Transcription analysis was performed on the ChIP-Seq dataset to highlight putative transcription factors that may functionally interact with ARID1A. We then interrogated the gene list from the transcriptomic dataset to understand how the activity of these transcription factors changed based on the expression of their target genes (Figure 4A). We identified increased activity of AP-1 family transcription factors to be associated with *Arid1a* loss (Figure 4B). The AP-1 transcription factor family is involved in critical cell processes such as differentiation and proliferation. Among the AP-1 subunits, *JunD* and *cFos* were both significantly upregulated following *Arid1a* loss by 1.5-fold and 3-fold respectively (Supplementary Table 3). We further investigate cFos and JUND protein levels in the murine tumours with varying *Arid1a* status. IHC staining confirmed dramatic upregulated nuclear cFos and JUND levels in *Pb-Cre;Pten^fl/fl^ Arid1a^fl/fl^* at 13-and 6.5-fold respectively, when compared to control *Pb-Cre;Pten^fl/fl^* and *Pb-Cre;Pten^fl/fl^ Arid1a^fl/+^* tumours (Figure 4C, Supplementary Figure 5A). This requirement for homozygous deletion of *Arid1a* in driving a pro-tumourigenic phenotype is reminiscent of the mouse survival data (Figure 2A).

**Figure 4:**
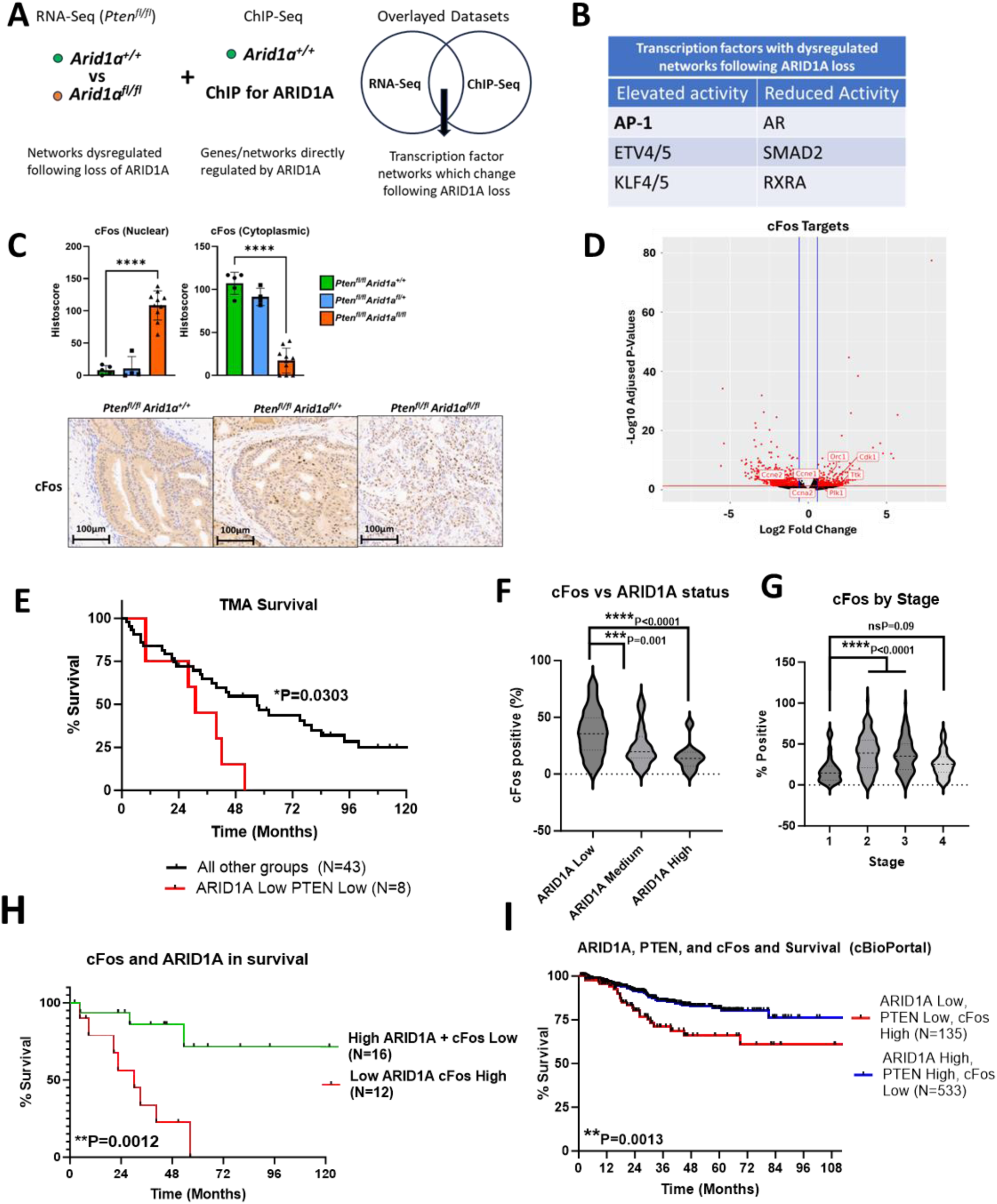
The status of ARID1A, AP-1 subunit cFos and PTEN is associated with patient survival. **A.** Schematic showing the overlaying of RNA-Seq and ChIP-Seq data to identify transcription factor networks regulated by ARID1A in mouse prostate tumours. **B.** Table showing transcription factors with significantly increased or decreased activity following ARID1A loss. **C.** Immunohistochemistry of indicated mouse prostate tissue stained and score for nuclear and cytoplasmic positivity of cFos including representative images. ****P<0.0001, ANOVA with Tukey’s analysis. Each data point is an individual mouse, error bars show SEM. **D.** Volcano plot showing up and downregulated cFos target genes (P<0.05, Fold Change >1.5) in *Pb-Cre;Pten^fl/fl^ Arid1a^fl/+^*(n=3) and *Pb-Cre;Pten^fl/fl^ Arid1a^fl/fl^* (n=3) cohorts. cFos target genes identified through dataset (https://maayanlab.cloud/Harmonizome/dataset/ENCODE+Transcription+Factor+Targets). **E.** Kaplan–Meier (log-rank) curve demonstrating survival of patient cohorts with different levels of ARID1A and PTEN as stained in a human PC tissue microarray. *P=0.0303; log-rank (Mantel-Cox) test. **F.** cFos positivity when compared to ARID1A status seen in Figure 1 from human prostate cancer tissue microarray. ***, P=0.001 ****P<0.0001; ANOVA with Tukey’s post hoc analysis. **G.** Histoscore for cFos staining of human tissue microarray by stage of prostate cancer (same as shown in Figure 1H), not significant P=0.09, ****P<0.0001; ANOVA with Tukey’s post hoc analysis. **H.** Kaplan–Meier (log-rank) curve demonstrating survival of patient cohorts with low ARID1A, High cFos compared to High ARID1A, Low cFos as stained in a human PC tissue microarray. **P=0.0012; log-rank (Mantel-Cox) test. **I.** Kaplan–Meier (log-rank) curve demonstrating survival of patients cohorts with different levels of *ARID1A*, *PTEN*, and *cFOS*. Patient data obtained from cBioPortal using studies of metastatic PC (SU2C/PCF Dream Team, Cell 2015), and primary PC (TCGA, Firehose Legacy). log-rank (Mantel-Cox) test ** p=0.0013.

By utilising publicly available cFos ChIP-Seq data from Kuonen *et al*, we interrogated how cFos target genes were changing in our mouse models (49). The observed increased expression and nuclear localisation of cFos are consistent with upregulated gene expression among known cFos target genes involved in cell cycle control such as *Cdk1*, *Cyclin E1*, *Cyclin E2*, and *A2* (Figure 4D). Interestingly, nuclear hormone receptors AR and RXRA were suggested to have reduced activity following *Arid1a* loss (Figure 4B), which is in line with previous observations that ARID1A can function as a transactivator of nuclear hormone receptors (10, 50).

We next investigated whether the status of ARID1A and PTEN in clinical tumours was associated with patient outcomes. Patients with tumours showing lower ARID1A and PTEN levels had a poorer survival compared to other patient groups combined (ARID1A low PTEN low median survival 31 months vs all other groups median survival 57 months *p=0.0303) (Figure 4E). We further explored the prognostic implications of altered cFos or JUND protein levels in our PC TMA. JUND did not correlate with tumour stage, ARID1A levels, or patient survival (Supplementary Figure 5B-E). Interestingly, cFos levels inversely correlated with ARID1A status (Figure 4F), while cFos levels increased as tumour stages increased from Stage 1 to 3, though not in Stage 4 tumours (Figure 4G). Importantly, combining ARID1A and cFos levels allows patient stratification into two prognostic groups, with low ARID1A/high cFos having a significantly reduced survival compared to high ARID1A/low cFos (Figure 4H). Interestingly this trend was not observed with cFos staining alone (Supplementary Figure 5F), suggesting that cFos is functionally related to ARID1A in driving prostate cancer progression. Finally, using publicly available clinical datasets from cBioPortal, tumours with low ARID1A, low PTEN, and high cFos were associated with a significantly poor survival outcome when compared to high ARID1A, high PTEN, and low cFos levels, corroborating the findings of our TMA analysis (Figure 4I).

## Discussion

Dysregulation of the epigenome is a hallmark of advanced cancers, with alterations in epigenetic regulators amongst the most frequently alterations found in in PC (51). Our Sleeping Beauty screen identified ARID1A as a candidate driver in PC (Figure 1). This finding was also reflected in clinical samples. In cBioPortal we observe frequent deletion of *ARID1A*; similarly, in our human TMA, reduced ARID1A protein levels were associated with less favourable patient survival outcome. The role of ARID1A in tumourigenesis appears diverse and context-dependent (9–11). This complex context-dependent role of ARID1A motivated us to investigate *Arid1a* using a GEMM system, where simultaneous loss of *Pten* and *Arid1a in vivo* produced aggressive and locally invasive prostate tumours (Figure 2).

Tumours from *Pb-Cre;Pten^fl/fl^ Arid1a^fl/fl^* mice have an interesting morphology, with reduced expression of luminal and increased expression of basal markers when compared to the tumours driven by homozygous *Pten* loss alone, suggestive of a less differentiated and proliferative phenotype (Figure 2C, D respectively). We further observed diminished and disorganised stroma in tumours driven by combined loss of *Pten* and *Arid1a*. To our knowledge, the *Pb-Cre;Pten^fl/fl^ Arid1a^fl/fl^* mouse model exhibited the most rapid tumour development to clinical endpoint of any published prostate cancer GEMM, with a hyperproliferative and locally invasive cancer (52). RNA-Seq identified that loss of *Arid1a* elevated cell cycle signalling (Figure 3). By overlaying the RNA-Seq and ChIP-Seq datasets, increased transcriptional activity of the AP-1 transcription factor family was suggested. This was consistent with our findings of enriched cell cycle-related genes being overrepresented and upregulated. Indeed, cFos, a key component of AP-1, when combined with ARID1A and PTEN, is found to be highly prognostic in a cohort of clinical prostate cancer (Figure 4).

A recent publication by Li *et al* 2022 also explored ARID1A in prostate cancer, and identified that loss of ARID1A can mediate immune evasion via a IKKβ/ARID1A/NF-κB axis (53). Immune evasion is expected to facilitate tumour initiation and metastasis while cell cycle elevation observed in our study will promote uncontrolled growth, as previously reported (12–15). Our study expands on the Li *et al* publication and further demonstrated that homozygous *Arid1a* deletion or deep loss of ARIDA expression is required for rapid prostate tumourigenesis: (1) Dramatic acceleration of tumourigenesis in *Pb-Cre;Pten^fl/fl^ Arid1a^fl/fl^* mice (Figure 2A), with solid tumour formation involving both the anterior and dorsolateral lobes while *Pb-Cre;Pten^fl/fl^* driven tumours are cystic and limited to the anterior lobes, (2) Tumour formation in the dorsolateral lobes of *Pb-Cre;Pten^fl/fl^ Arid1a^fl/fl^*mice originated from the successful progression of PIN lesions in the dorsolateral lobes of *Pb-Cre;Pten^fl/fl^* mice into tumours (Supplementary Table 2), (3) Upregulated cFos and JUND expression in tumours from *Pb-Cre;Pten^fl/fl^ Arid1a^fl/fl^*mice, (4) Significant increased colony forming ability in DU145 ARID1A KO2 and KO4 clones with negligible ARID1A expression (Supplementary Figure 4), (5) Poor patient survival outcome being associated with low ARID1A and PTEN expression, and low ARID1A and high cFos expression in our TMA PC cohort (Figure 4E, 4H respectively), and (6) Association between reduced ARID1a and increased cFos expression (Figure 4F) and the poor patient outcome for tumours with high cFos, low PTEN and low ARID1A expression (Figure 4I). It is worth noting that our observation of reduced and disorganised collagen expression in tumours from *Pb-Cre;Pten^fl/fl^ Arid1a^fl/fl^* mice is consistent with the model whereby a collagen-poor stroma results in enriched tumour-suppressive cytokines and leads to undifferentiated and invasive pancreatic cancer with shorted patient survival (54). The focus of future study will help determine the interplay between cancer and immune cells within the tumour microenvironment.

Based on the publicly available datasets in cBioPortal, shallow, rather than deep, *ARID1A* deletions are documented, implicating additional genetic and epigenetic events in order to accelerate tumourigenesis to the level observed in our *Pb-Cre;Pten^fl/fl^ Arid1a^fl/fl^* mouse cohort. Future research is warranted to fully defined molecular events that would interact with shallow loss of *ARID1A* in clinical tumours. Previous studies have also demonstrated the *Pten*-deficient murine models stabilise BRG1 allowing the SWI/SNF to mediate oncogenic remodelling in a BRG1-dependent manner(55). This would suggest an ARID1B-BRG1 BAF complex may represent a particularly potent ‘onco-BAF’ complex in PTEN-deficient PC, in particular with loss of ARID1A. Importantly, this subtype of PC may be targetable through exploiting their defective DNA-damage response as has been demonstrated in other ARID1A-mutant cancers (56, 57). This can include targeting DNA-damage response machinery, such as through PARP, ATM, or ATR inhibition as single agents or as radiosensitisers (56, 58–61). Alternatively, BRM/BRG1 PROTACS may be of efficacy in cancers with mutated BAF components (62).

## Conclusions

Homozygous *Arid1a* loss dramatically accelerates prostate tumourigenesis, resulting in hyper-proliferative and undifferentiated tumours with a reduced and disorganised stroma. *Arid1a* loss mediated tumour formation in the mouse involved both the anterior and dorsoateral lobes, a key distinction from *Pten*-loss driven tumours which tend to be limited to the anterior lobes. Finally, the status of PTEN, ARID1A and cFos, as an ARID1A downstream effector, is associated with patient survival outcome.

## Acknowledgements

We wish to thank the core facilities, biological services unit, and molecular services at the CRUK Scotland Institute.

## Abbreviations

BAF: BRM/BRG1 associated factors
BRG1: Brahma-related gene 1
BRM: Brahma
ChIP: Chromatin Immunoprecpitation
ER: Oestrogen receptor
Fl: Flox
HGPIN: High grade intraepithelial neoplasm
IHC: Immunohistochemistry
KO: Knockout
LGPIN: Low grade intraepithelial neoplasm
Pb-Cre: Probasin Cre-recombinase
PC: Prostate Cancer
PCA: Principal component analysis
rEGF: recombinant epidermal growth factor
SB: Sleeping Beauty
SCM: Standard Culture Medium
SWI/SNF: Switch-induced/sucrose non-fermentable
TMA: Tissue microarray

## Declarations

## Ethical Approval

Animal experiments were carried out in line with the Animals (Scientific Procedures) Act 1986 and the EU Directive of 2010 sanctioned by Local Ethical Review Process (University of Glasgow). Mice were maintained on a mixed strain background at the Cancer Research UK Scotland Institute under project licence authority (70/8645 and P5EE22AEE to Professor Hing Leung).

## Author Contributions

Conception and design: AH and IA. Supervision: KB, HL and IA. Development of methodology: AH, RS, LG and RH. Acquisition of data: AH, LG, AT. Analysis and interpretation of data: AH, RS, RV, LW, IA. Writing, review and/or revision of the manuscript: AH, KB, HL, IA.

## Funding Information

CRUK Scotland Institute (A31287)

CRUK core funding for KB (A29799)

CRUK Scotland Centre (CTRQQR-2021\100006)

CRUK Clinician Scientist Fellowship (19661) to IA.

Prostate Cancer Foundation PCF#18CHAL11 to RH.

## Availability of data and materials

Data will be made available prior to publication.

---

**Supplementary Figure 1:**
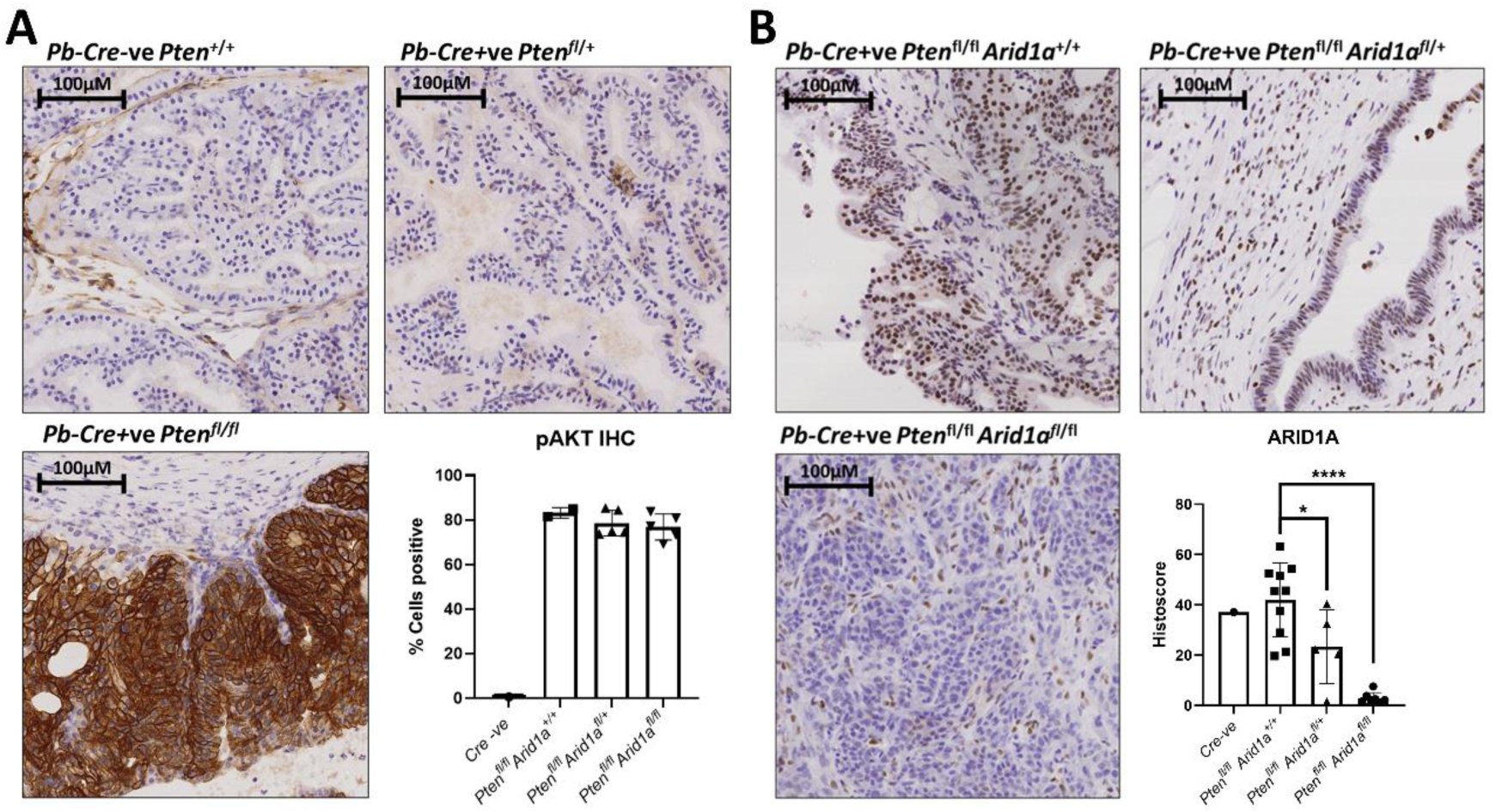
Immunohistochemistry for phospho-serine 474 AKT and ARID1A. **A.** Representative staining of prostate stained for phospho-serine 474 AKT and quantification of stain-positive cells. *Pb-Cre:Pten^+/+^* shown at 6 months timepoint, *Pb-Cre:Pten^fl/+^*shown at 6 month timepoint, *Pb-Cre:Pten^fl/fl^* shown at 9 month clinical endpoint. Quantification of pAKT IHC in Cre-ve compared to *Pb-Cre:Pten^fl/fl^* with various *Arid1a* status shows ARID1A loss does not impact pAKT levels. Each data point is an individual mouse, error bars show SEM. **B.** Representative staining of endpoint prostate tumour stained for ARID1A and quantification of histoscore in indicated genotypes. Each data point represents an individual mouse, error bars show SEM.*P=0.0373, ****P<0.0001 ANOVA with Tukey’s analysis.

**Supplementary Figure 2:**
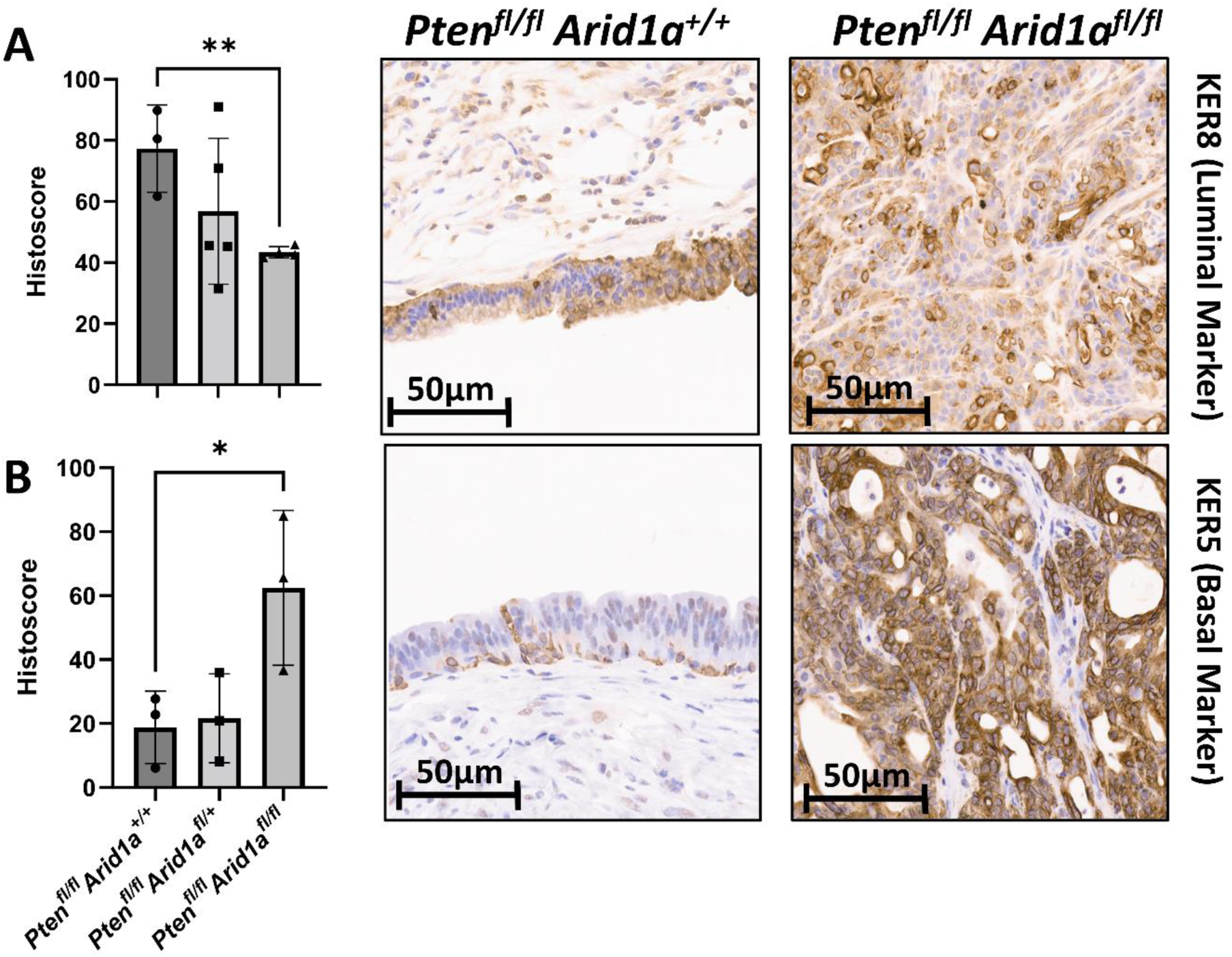
IHC for basal vs luminal markers following loss of ARID1A. **A.** Representative staining of Keratin 8 in epithelial compartment of indicated genotypes. Quantification of histoscore in indicated genotype, **P=0.0047 Keratin 8; tested by ANOVA with Tukey’s post hoc analysis. Each data point is an individual tumour. **B** Representative staining of Keratin 5 in the indicated genotypes. Quantification of histoscore in epithelial compartment of indicated genotype, *P=0.048 Keratin 5; tested by ANOVA with Tukey’s post hoc analysis.

**Supplementary Figure 3:**
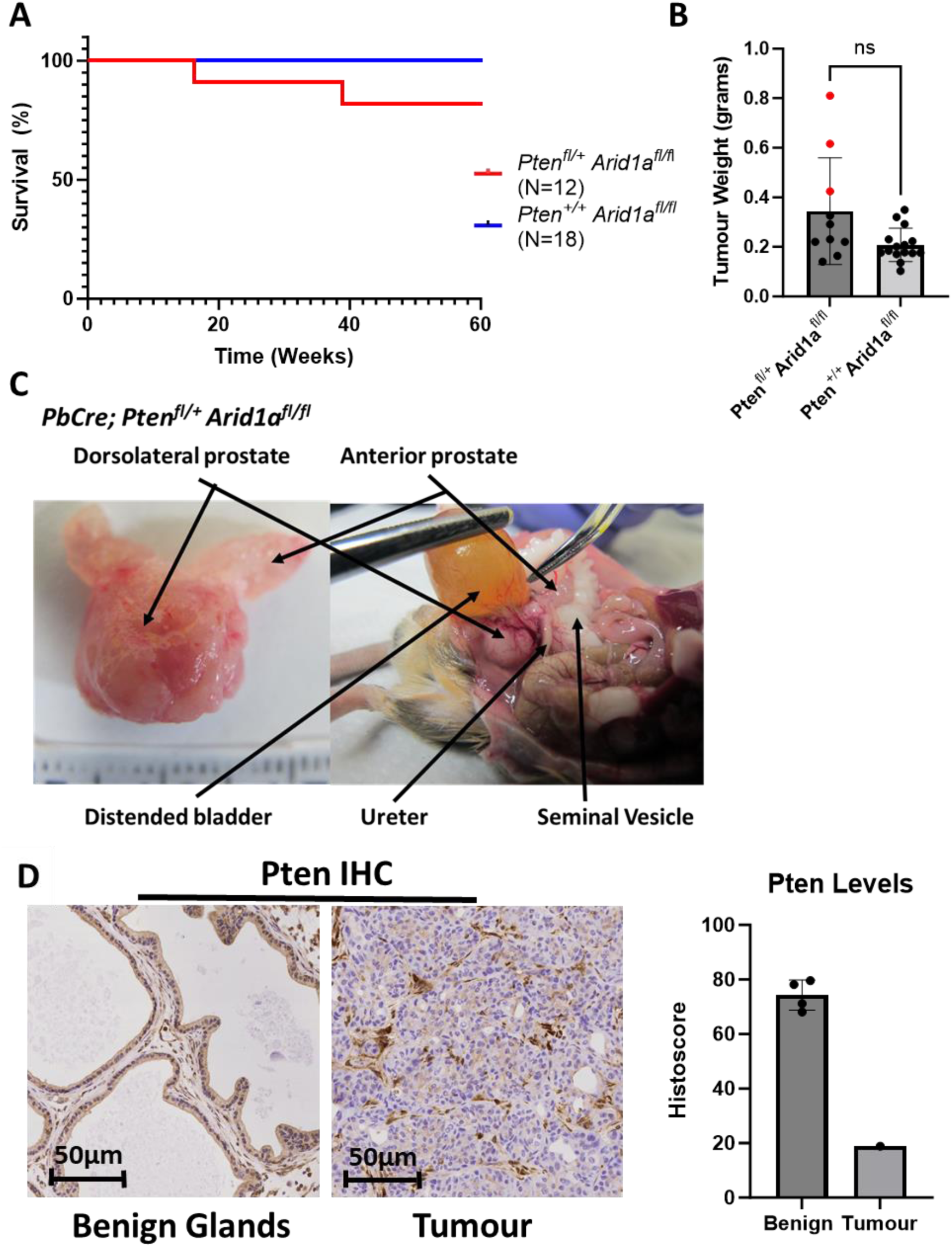
*Pten* loss is a prerequisite for *Arid1a* loss mediated tumorigenesis. **A.** Kaplan–Meier (log-rank) curve demonstrating survival of *Pb-Cre;Pten^fl/+^Arid1a^fl/fl^*(n=17) and *Pb-Cre;Pten^+/+^ Arid1a^fl/fl^* (n=18) male mice, not significant; log-rank (Mantel-Cox) test. **B.** Weight in grams of prostate/tumours harvested from male mice taken at clinical or ageing endpoint at 60 weeks of *Pb-Cre;Pten^fl/+^Arid1a^fl/fl^*(n=12 and *Pb-Cre;Pten^+/+^ Arid1a^fl/fl^*(n=18), not significant P=0.083, Mann-Whitney. Each data point represents and individual mouse, those in red reached clinical endpoint due to tumour burden. **C.** *Ex vivo* and *in situ* prostate tumour of *Pb-Cre;Pten^fl/+^Arid1a^fl/fl^* genotype with spontaneous development of dorsolateral tumour. with key structures labelled. **D.** Staining of endpoint dorsolateral prostate tumour of a *Pb-Cre;Pten^fl/+^Arid1a^fl/fl^* mouse for PTEN and ARID1A respectively with key structures labelled.

**Supplementary Figure 4:**
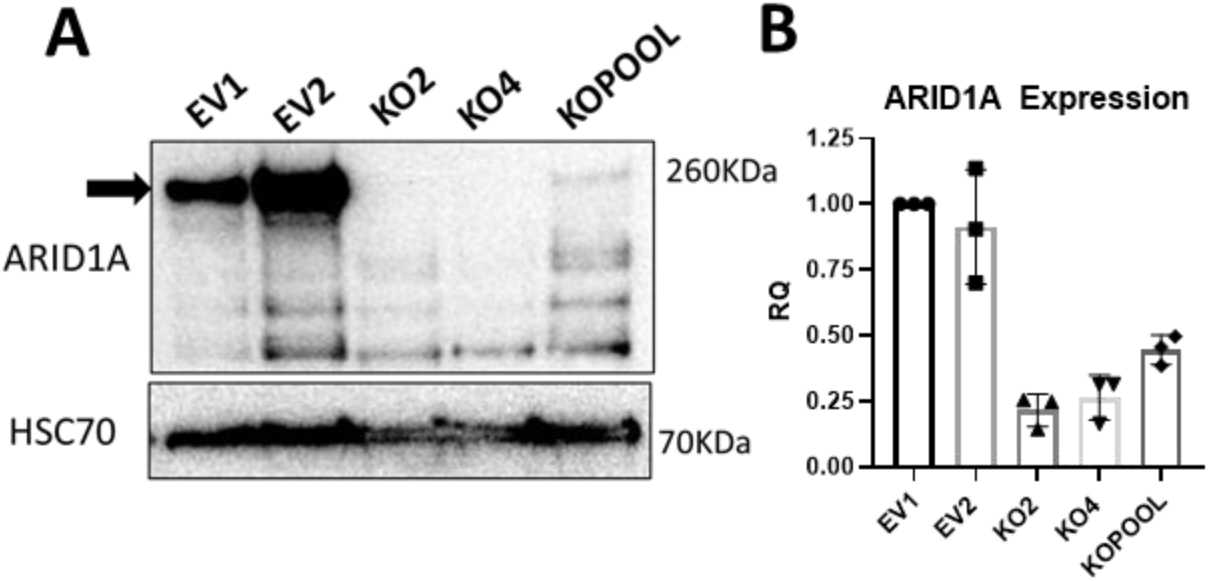
Validation of CRISPR-Cas9 Knockout of ARID1A in DU145 cells. **A.** Immunoblotting of protein from DU145 empty vector (EV) 1 and 2 clones, and ARID1A knockout (KO) clones 2, 4, and pool for ARID1A (Molecular weight = 260 kDa). HSC70 was used as loading control. Representative blot of three experimental replicates. **B.** Quantitative RT-PCR comparing relative quantification (RQ) of *ARID1A* expression in DU145 EV clones compared to *ARID1A* KO clones. Each point represents an experimental replicate each made up of three technical replicates, error bars showing SEM.

**Supplementary Figure 5:**
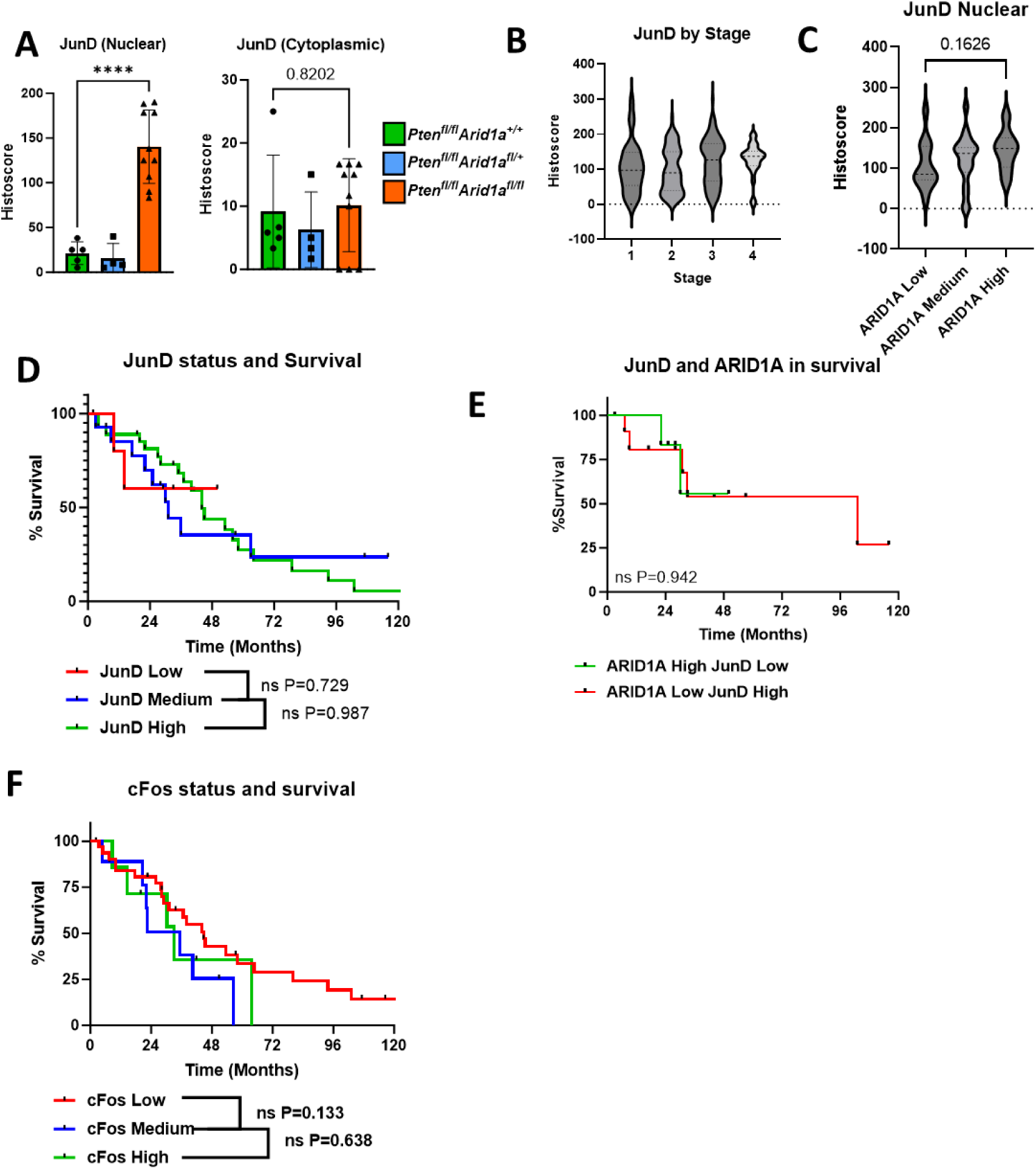
A. JunD status does not predict prostate cancer survival. Immunohistochemistry of indicated mouse tissue (as in Figure 5) stained and scored for nuclear and cytoplasmic positivity of JunD ****P<0.0001. Each data point is an individual mouse. **B.** Histoscore for nuclear JunD staining of human tissue microarray by stage of prostate cancer (same as in Figure 1 Iand 5). **C.** JunD positivity when compared to ARID1A status from human prostate cancer tissue microarray. Not significant, P=0.1626; ANOVA with Tukey’s analysis. **D.** Kaplan–Meier (log-rank) curve demonstrating survival of patient cohorts with different levels of JunD. JunD Low (N=5); JunD Medium (N=14); JunD High (N=28). Not significant; log-rank (Mantel-Cox) test. **E.** Kaplan–Meier (log-rank) curve demonstrating survival of patient cohorts comparing ARID1A low JunD high (N=12) vs ARID1A high JunD low (N=6). Not significant; log-rank (Mantel-Cox) test. **F.** Kaplan–Meier (log-rank) curve demonstrating survival of patient cohorts with different levels of cFos stained in the TMA. cFos Low (N=31), cFos Medium (N=10), cFos High (N=7). Not significant; log-rank (Mantel-Cox) test.

**Supplementary Table 1.**
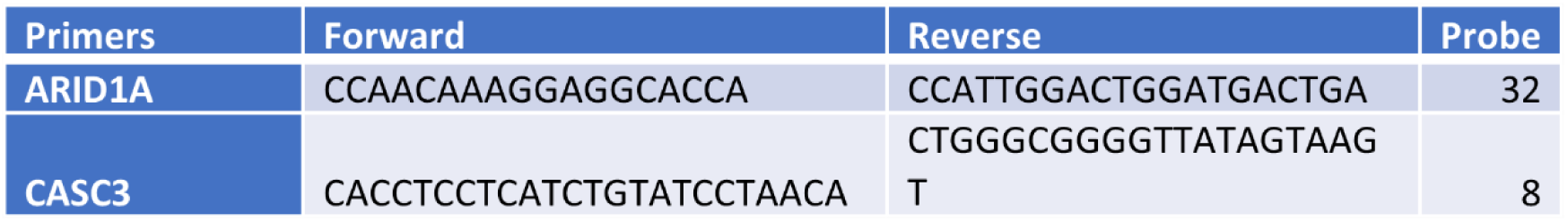
Primers QPCR primer target, forward and reverse sequences, and Roche Probe used for detection with Taqman Reagent.

**Supplementary Table 2.** (A) Proportions of mice which developed prostate cancer of anterior or dorsolateral prostate neoplasia as graded as resembling normal tissue, prostate intraepithelial neoplasm (either low grade prostate intraepithelial neoplasm (LGPIN) or high grade prostate intraepithelial neoplasm (HGPIN)), adenocarcinoma, and having reached clinical endpoint due to prostatic tumour. (n/N, number of mice affected/total number of mice in the group). (B) Extract of data from panel A to highlight the incidence of tumour formation in the anterior and dorsolateral lobes of the mice with homozygous *Pten* deletion and varying status of *Arid1a*. (n/N, number of mice affected/total number of mice in the group).

| Genotype | Normal (n/N) | LGPIN (n/N) | HGPIN (n/N) | Adenocarcinoma (n/N) | Tumour endpoint (n/N) |
| --- | --- | --- | --- | --- | --- |
| **Anterior** |  |  |  |  |  |
| *Pten*<sup>+/+</sup>*Arid1a*<sup>fl/fl</sup> | 15/17 | 2/17 |  |  |  |
| *Pten*<sup>fl/+</sup>*Arid1a*<sup>fl/fl</sup> | 9/12 | 2/12 |  | 1/12 |  |
| *Pten*<sup>fl/fl</sup> *Arid1a*<sup>+/+</sup> |  |  |  |  | 10/10 Cystic |
| *Pten*<sup>fl/fl</sup> *Arid1a*<sup>fl/+</sup> |  |  |  |  | 19/19 Cystic |
| *Pten*<sup>fl/fl</sup> *Arid1a*<sup>fl/fl</sup> |  |  |  |  | 8/8 Solid |
| **Dorsolateral** |  |  |  |  |  |
| *Pten*<sup>+/+</sup>*Arid1a*<sup>fl/fl</sup> | 10/17 | 7/17 |  |  |  |
| *Pten*<sup>fl/+</sup>*Arid1a*<sup>fl/fl</sup> | 4/12 | 2/12 | 1/12 | 2/12 | 3/12 |
| *Pten*<sup>fl/fl</sup> *Arid1a*<sup>+/+</sup> |  |  | 10/10 |  |  |
| *Pten*<sup>fl/fl</sup> *Arid1a*<sup>fl/+</sup> |  |  | 19/19 |  |  |
| *Pten*<sup>fl/fl</sup> *Arid1a*<sup>fl/fl</sup> |  |  |  |  | 8/8 Solid |

**Supplementary Table 2.** (A) Proportions of mice which developed prostate cancer of anterior or dorsolateral prostate neoplasia as graded as resembling normal tissue, prostate intraepithelial neoplasm (either low grade prostate intraepithelial neoplasm (LGPIN) or high grade prostate intraepithelial neoplasm (HGPIN)), adenocarcinoma, and having reached clinical endpoint due to prostatic tumour.
|  | *Pten*<sup>fl/fl</sup> *Arid1a*<sup>+/+</sup><br>n/N | *Pten*<sup>fl/fl</sup> *Arid1a*<sup>fl/+</sup><br>n/N | *Pten*<sup>fl/fl</sup> *Arid1a*<sup>fl/fl</sup><br>n/N |
| --- | --- | --- | --- |
| **Anterior** | Cystic Tumours 10/10 | Cystic Tumours 19/19 | Solid Tumours 8/8 |
| **Dorsolateral** | No tumour | No tumour | Solid Tumours 8/8 |

**Supplementary Table 3.**
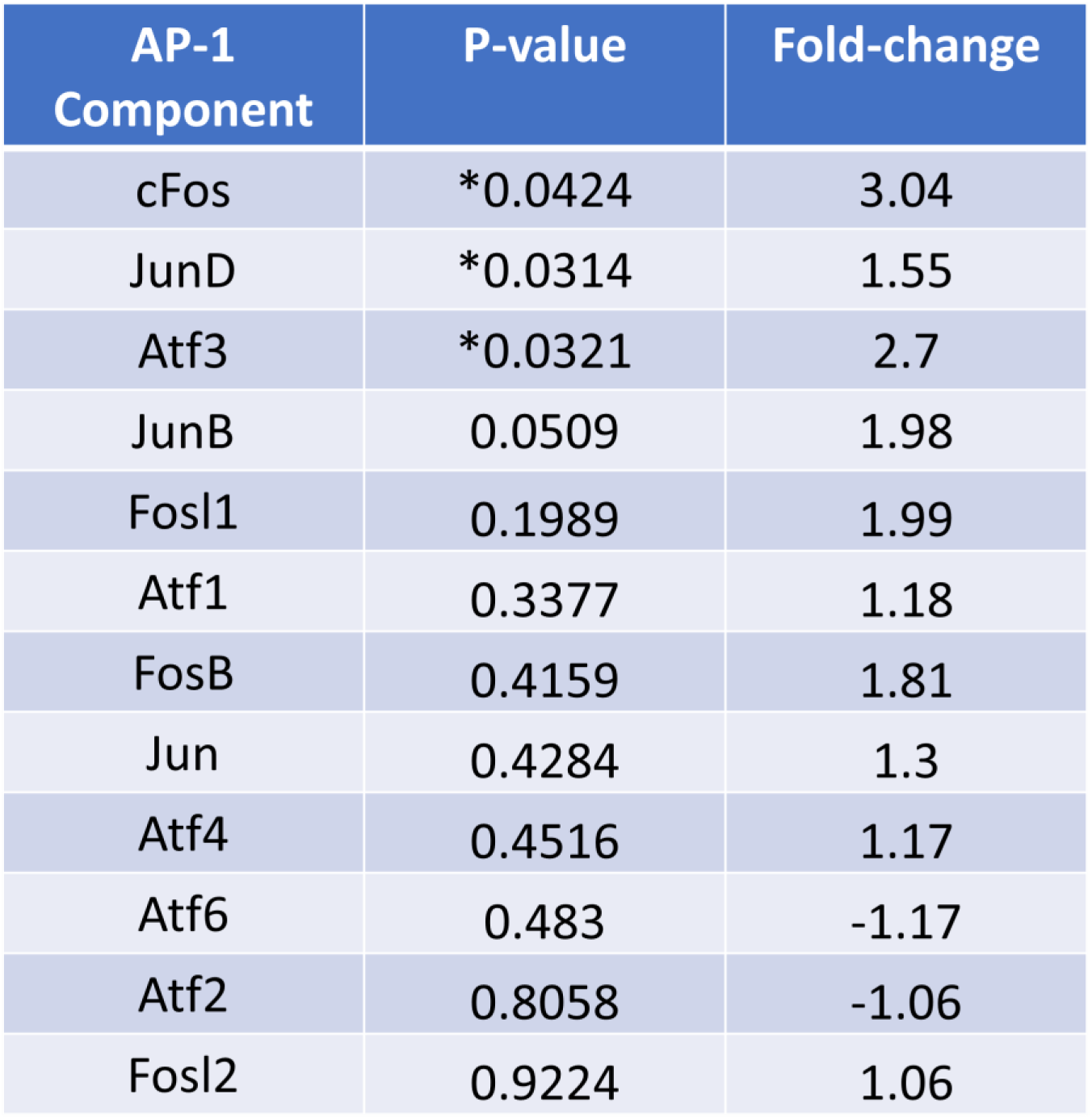
AP-1 components showing fold change and degree of significance comparing *Pb-Cre;Pten^fl/fl^ Arid1a^fl/+^* (n=3) and *Pb-Cre;Pten^fl/fl^ Arid1a^fl/fl^* (n=3) cohorts.

| AP-1 Component | P-value | Fold-change |
| --- | --- | --- |
| cFos | *0.0424 | 3.04 |
| JunD | *0.0314 | 1.55 |
| Atf3 | *0.0321 | 2.7 |
| JunB | 0.0509 | 1.98 |
| Fosl1 | 0.1989 | 1.99 |
| Atf1 | 0.3377 | 1.18 |
| FosB | 0.4159 | 1.81 |
| Jun | 0.4284 | 1.3 |
| Atf4 | 0.4516 | 1.17 |
| Atf6 | 0.483 | -1.17 |
| Atf2 | 0.8058 | -1.06 |
| Fosl2 | 0.9224 | 1.06 |

**Supplementary Table 4.** GSEA performed comparing mouse cohorts *Pb-Cre;Pten^fl/fl^ Arid1a^fl/+^* (n=3) and *Pb-Cre;Pten^fl/fl^ Arid1a^fl/fl^* (n=3), and DU145 EV1 (n=5) vs ARID1A KO2 (n=5) cohorts. Indicated hallmark pathway shown along with normalised enrichment scores (NES), and P-values.

| Hallmark Pathway | Mouse: NES;P-value | DU145: NES;P-value |
| --- | --- | --- |
| TGF_BETA_SIGNALING | 1.00; 0.521 | 2.21; 0.000 |
| TNFA_SIGNALING_VIA_NFKB | 0.88; 0.791 | 2.57; 0.000 |
| INTERFERON_GAMMA_RESPONSE | 0.892; 0.907 | 1.78; 0.000 |
| INFLAMMATORY_RESPONSE | 0.891; 1.000 | 2.02;0.000 |
| IL6_JAK_STAT3_SIGNALING | 0.891; 0.988 | 1.82;0.000 |

## References

1. UK CR. Prostate Cancer Statistics https://www.cancerresearchuk.org/health-professional/cancer-statistics/statistics-by-cancer-type/prostate-cancer [

2. Ahmad I, Mui E, Galbraith L, Patel R, Tan EH, Salji M, et al. Sleeping Beauty screen reveals Pparg activation in metastatic prostate cancer. Proceedings of the National Academy of Sciences. 2016;113(29):8290.

3. Tibbo AJ, Hartley A, Vasan R, Shaw R, Galbraith L, Mui E, et al. MBTPS2 acts as a regulator of lipogenesis and cholesterol synthesis through SREBP signalling in prostate cancer. Br J Cancer. 2023;128(11):1991–9.

4. Tang L, Nogales E, Ciferri C. Structure and function of SWI/SNF chromatin remodeling complexes and mechanistic implications for transcription. Progress in biophysics and molecular biology. 2010;102(2-3):122–8.

5. Yegnasubramanian S, De Marzo AM, Nelson WG. Prostate Cancer Epigenetics: From Basic Mechanisms to Clinical Implications. Cold Spring Harb Perspect Med. 2019;9(4).

6. Flowers S, Nagl NG, Jr., Beck GR, Jr., Moran E. Antagonistic roles for BRM and BRG1 SWI/SNF complexes in differentiation. The Journal of biological chemistry. 2009;284(15):10067–75.

7. Wu JN, Roberts CW. ARID1A mutations in cancer: another epigenetic tumor suppressor? Cancer discovery. 2013;3(1):35–43.

8. Wu RC, Wang TL, Shih Ie M. The emerging roles of ARID1A in tumor suppression. Cancer biology & therapy. 2014;15(6):655–64.

9. Sen M, Wang X, Hamdan FH, Rapp J, Eggert J, Kosinsky RL, et al. ARID1A facilitates KRAS signaling-regulated enhancer activity in an AP1-dependent manner in colorectal cancer cells. Clinical Epigenetics. 2019;11(1):92.

10. Xu G, Chhangawala S, Cocco E, Razavi P, Cai Y, Otto JE, et al. ARID1A determines luminal identity and therapeutic response in estrogen-receptor-positive breast cancer. Nature genetics. 2020;52(2):198–207.

11. Hiramatsu Y, Fukuda A, Ogawa S, Goto N, Ikuta K, Tsuda M, et al. Arid1a is essential for intestinal stem cells through Sox9 regulation. Proceedings of the National Academy of Sciences. 2019;116(5):1704–13.

12. Wu C, Lyu J, Yang EJ, Liu Y, Zhang B, Shim JS. Targeting AURKA-CDC25C axis to induce synthetic lethality in ARID1A-deficient colorectal cancer cells. Nature Communications. 2018;9(1):3212.

13. Shen J, Peng Y, Wei L, Zhang W, Yang L, Lan L, et al. ARID1A Deficiency Impairs the DNA Damage Checkpoint and Sensitizes Cells to PARP Inhibitors. Cancer discovery. 2015;5(7):752–67.

14. Park Y, Chui MH, Suryo Rahmanto Y, Yu Z-C, Shamanna RA, Bellani MA, et al. Loss of ARID1A in tumor cells renders selective vulnerability to combined ionizing radiation and PARP inhibitor therapy. Clinical Cancer Research. 2019:clincanres.4222.2019.

15. Jones S, Wang T-L, Shih I-M, Mao T-L, Nakayama K, Roden R, et al. Frequent Mutations of Chromatin Remodeling Gene *ARID1A* in Ovarian Clear Cell Carcinoma. Science. 2010;330(6001):228–31.

16. Sandoval GJ, Pulice JL, Pakula H, Schenone M, Takeda DY, Pop M, et al. Binding of TMPRSS2-ERG to BAF Chromatin Remodeling Complexes Mediates Prostate Oncogenesis. Molecular cell. 2018;71(4):554–66.e7.

17. Dufour CR, Scholtes C, Yan M, Chen Y, Han L, Li T, et al. The mTOR chromatin-bound interactome in prostate cancer. Cell Rep. 2022;38(12):110534.

18. Wu X, Wu J, Huang J, Powell WC, Zhang J, Matusik RJ, et al. Generation of a prostate epithelial cell-specific Cre transgenic mouse model for tissue-specific gene ablation. Mech Dev. 2001;101(1-2):61–9.

19. Suzuki A, Yamaguchi MT, Ohteki T, Sasaki T, Kaisho T, Kimura Y, et al. T cell-specific loss of Pten leads to defects in central and peripheral tolerance. Immunity. 2001;14(5):523–34.

20. March HN, Rust AG, Wright NA, ten Hoeve J, de Ridder J, Eldridge M, et al. Insertional mutagenesis identifies multiple networks of cooperating genes driving intestinal tumorigenesis. Nature genetics. 2011;43(12):1202–9.

21. Gao X, Tate P, Hu P, Tjian R, Skarnes WC, Wang Z. ES cell pluripotency and germ-layer formation require the SWI/SNF chromatin remodeling component BAF250a. Proc Natl Acad Sci U S A. 2008;105(18):6656–61.

22. Ahmad I, Patel R, Liu Y, Singh LB, Taketo MM, Wu XR, et al. Ras mutation cooperates with β-catenin activation to drive bladder tumourigenesis. Cell Death Dis. 2011;2(3):e124.

23. Bioinformatics ASB. FastQC A Quality Control tool for High Throughput Sequence Data [Available from: https://www.bioinformatics.babraham.ac.uk/projects/fastqc.

24. Chen S, Zhou Y, Chen Y, Gu J. fastp: an ultra-fast all-in-one FASTQ preprocessor. Bioinformatics. 2018;34(17):i884–i90.

25. Wingett SW, Andrews S. FastQ Screen: A tool for multi-genome mapping and quality control. F1000Res. 2018;7:1338.

26. Kim D, Paggi JM, Park C, Bennett C, Salzberg SL. Graph-based genome alignment and genotyping with HISAT2 and HISAT-genotype. Nat Biotechnol. 2019;37(8):907–15.

27. Li H, Handsaker B, Wysoker A, Fennell T, Ruan J, Homer N, et al. The Sequence Alignment/Map format and SAMtools. Bioinformatics. 2009;25(16):2078–9.

28. Liao Y, Smyth GK, Shi W. featureCounts: an efficient general purpose program for assigning sequence reads to genomic features. Bioinformatics. 2014;30(7):923–30.

29. Team RC. R: A Language and Environment for Statistical Computing [Available from: https://www.R-project.org/.

30. Huber W, Carey VJ, Gentleman R, Anders S, Carlson M, Carvalho BS, et al. Orchestrating high-throughput genomic analysis with Bioconductor. Nat Methods. 2015;12(2):115–21.

31. Love MI, Huber W, Anders S. Moderated estimation of fold change and dispersion for RNA-seq data with DESeq2. Genome Biol. 2014;15(12):550.

32. Stephens M. False discovery rates: a new deal. Biostatistics. 2017;18(2):275–94.

33. Cunningham F, Achuthan P, Akanni W, Allen J, Amode MR, Armean IM, et al. Ensembl 2019. Nucleic Acids Res. 2019;47(D1):D745–d51.

34. Howe KL, Achuthan P, Allen J, Allen J, Alvarez-Jarreta J, Amode MR, et al. Ensembl 2021. Nucleic Acids Res. 2021;49(D1):D884–d91.

35. Raudvere U, Kolberg L, Kuzmin I, Arak T, Adler P, Peterson H, et al. g:Profiler: a web server for functional enrichment analysis and conversions of gene lists (2019 update). Nucleic Acids Research. 2019;47(W1):W191–W8.

36. Subramanian A, Tamayo P, Mootha VK, Mukherjee S, Ebert BL, Gillette MA, et al. Gene set enrichment analysis: a knowledge-based approach for interpreting genome-wide expression profiles. Proc Natl Acad Sci U S A. 2005;102(43):15545–50.

37. Ewels PA, Peltzer A, Fillinger S, Patel H, Alneberg J, Wilm A, et al. The nf-core framework for community-curated bioinformatics pipelines. Nat Biotechnol. 2020;38(3):276–8.

38. Fornes O, Castro-Mondragon JA, Khan A, van der Lee R, Zhang X, Richmond PA, et al. JASPAR 2020: update of the open-access database of transcription factor binding profiles. Nucleic Acids Res. 2020;48(D1):D87–d92.

39. Quinlan AR, Hall IM. BEDTools: a flexible suite of utilities for comparing genomic features. Bioinformatics. 2010;26(6):841–2.

40. Wang Z, Civelek M, Miller CL, Sheffield NC, Guertin MJ, Zang C. BART: a transcription factor prediction tool with query gene sets or epigenomic profiles. Bioinformatics. 2018;34(16):2867–9.

41. Wickham H, Averick M, Bryan J, Chang W, McGowan L, François R, et al. Welcome to the Tidyverse. Journal of Open Source Software. 2019;4:1686.

42. Ewels P, Magnusson M, Lundin S, Käller M. MultiQC: summarize analysis results for multiple tools and samples in a single report. Bioinformatics. 2016;32(19):3047–8.

43. T. K. Jupyter Notebooks—a publishing format for reproducible computational workflows IOS Press. 2016.

44. Team R. Rstudio: Integrated Development Environment for R 2019.

45. Rooney N, Mason SM, McDonald L, Däbritz JHM, Campbell KJ, Hedley A, et al. RUNX1 Is a Driver of Renal Cell Carcinoma Correlating with Clinical Outcome. Cancer Research. 2020;80(11):2325–39.

46. Ahmad I, Patel R, Singh LB, Nixon C, Seywright M, Barnetson RJ, et al. HER2 overcomes PTEN (loss)-induced senescence to cause aggressive prostate cancer. Proc Natl Acad Sci U S A. 2011;108(39):16392–7.

47. Wang S, Gao J, Lei Q, Rozengurt N, Pritchard C, Jiao J, et al. Prostate-specific deletion of the murine Pten tumor suppressor gene leads to metastatic prostate cancer. Cancer Cell. 2003;4(3):209–21.

48. Raab JR, Resnick S, Magnuson T. Genome-Wide Transcriptional Regulation Mediated by Biochemically Distinct SWI/SNF Complexes. PLOS Genetics. 2016;11(12):e1005748.

49. Kuonen F, Li NY, Haensel D, Patel T, Gaddam S, Yerly L, et al. c-FOS drives reversible basal to squamous cell carcinoma transition. Cell Reports. 2021;37(1):109774.

50. Launonen K-M, Paakinaho V, Sigismondo G, Malinen M, Sironen R, Hartikainen JM, et al. Chromatin-directed proteomics-identified network of endogenous androgen receptor in prostate cancer cells. Oncogene. 2021;40(27):4567–79.

51. Hartley A, Leung HY, Ahmad I. Targeting the BAF complex in advanced prostate cancer. Expert Opin Drug Discov. 2021;16(2):173–81.

52. Arriaga JM, Abate-Shen C. Genetically Engineered Mouse Models of Prostate Cancer in the Postgenomic Era. Cold Spring Harb Perspect Med. 2019;9(2).

53. Li N, Liu Q, Han Y, Pei S, Cheng B, Xu J, et al. ARID1A loss induces polymorphonuclear myeloid-derived suppressor cell chemotaxis and promotes prostate cancer progression. Nature Communications. 2022;13(1):7281.

54. Chen Y, Kim J, Yang S, Wang H, Wu CJ, Sugimoto H, et al. Type I collagen deletion in αSMA(+) myofibroblasts augments immune suppression and accelerates progression of pancreatic cancer. Cancer Cell. 2021;39(4):548–65.e6.

55. Ding Y, Li N, Dong B, Guo W, Wei H, Chen Q, et al. Chromatin remodeling ATPase BRG1 and PTEN are synthetic lethal in prostate cancer. J Clin Invest. 2019;129(2):759–73.

56. Shen J, Peng Y, Wei L, Zhang W, Yang L, Lan L, et al. ARID1A Deficiency Impairs the DNA Damage Checkpoint and Sensitizes Cells to PARP Inhibitors. Cancer discovery. 2015;5(7):752–67.

57. Mandal J, Mandal P, Wang T-L, Shih I-M. Treating ARID1A mutated cancers by harnessing synthetic lethality and DNA damage response. Journal of Biomedical Science. 2022;29(1):71.

58. Williamson CT, Miller R, Pemberton HN, Jones SE, Campbell J, Konde A, et al. ATR inhibitors as a synthetic lethal therapy for tumours deficient in ARID1A. Nature Communications. 2016;7(1):13837.

59. Wang L, Yang L, Wang C, Zhao W, Ju Z, Zhang W, et al. Inhibition of the ATM/Chk2 axis promotes cGAS/STING signaling in ARID1A-deficient tumors. J Clin Invest. 2020;130(11):5951–66.

60. Yang L, Yang G, Ding Y, Dai Y, Xu S, Guo Q, et al. Inhibition of PI3K/AKT Signaling Pathway Radiosensitizes Pancreatic Cancer Cells with ARID1A Deficiency in Vitro. J Cancer. 2018;9(5):890–900.

61. Niedermaier B, Sak A, Zernickel E, Xu S, Groneberg M, Stuschke M. Targeting ARID1A-mutant colorectal cancer: depletion of ARID1B increases radiosensitivity and modulates DNA damage response. Scientific Reports. 2019;9(1):18207.

62. Xiao L, Parolia A, Qiao Y, Bawa P, Eyunni S, Mannan R, et al. Targeting SWI/SNF ATPases in enhancer-addicted prostate cancer. Nature. 2022;601(7893):434–9.

